# A scaling theory of trait evolution

**DOI:** 10.1101/2025.11.18.689082

**Authors:** W. Bryan Jennings

## Abstract

A scaling model named the “geometric-similarity-first model” is developed to explain allometric trait divergence over recent evolutionary time. In the model’s first step, traits undergo geometric scaling with a population’s change in body size. In step 2, directional natural selection re-optimizes trait shapes such that traits showing positive ontogenetic allometry undergo positive evolutionary allometric scaling while traits that exhibit negative ontogenetic allometry go through negative evolutionary allometric scaling. Five predictions of the model were tested using morphological data for three locomotor-relevant traits in pygopodid lizards. The dataset, which was based on 1,756 museum specimens representing 31 species, supported all of these predictions. An implication of these results is that geometric scaling, driven by natural or sexual selection, may be a mechanism for peak shifts on an adaptive landscape. Given the ubiquity of body size variation in nature, this hypothetical process, termed “niche scaling,” may be important to ecological diversification. Applications of the model to some other well-studied species are discussed.

## INTRODUCTION

Since Darwin, evolutionary biologists have attempted to understand the mechanisms of functional trait divergence because this process played a major role in generating the diversity of life (Darwin 1859; Lack 1947; Simpson 1944, 1953; Grant 1986; Schluter 2000; Grant and Grant 2008; Losos 2009; Gillespie et al. 2020; Tobias et al. 2021; Grant and Grant 2024). Although natural selection’s importance as a driver of trait divergences has been shown in many studies, aspects of this process—especially during the initial stage of trait divergence—remain obscure (Schluter 2000; Stroud and Losos 2020). The problem of initial divergence is far-reaching in evolutionary biology. For example, we still know little about how natural and sexual selection drive peak shifts on adaptive landscapes (Simpson 1944, 1953; Schluter 2000; Bonduriansky 2011).

Analyses of paired allometric growth trajectories between populations and closely related species can reveal distinctive trait divergence patterns, each with its own biomechanical and macroevolutionary implications (Kurtén 1954, 1955; Gould 1966, 1971; Boag 1984; Klingenberg 1998; Sanger et al. 2012; Stroud et al. 2023). For example, geometric similarity (hereafter GS) in traits can be manifested as two dramatically different patterns on log-log plots depending on whether a trait shows positive or negative ontogenetic (or static) allometry (hereafter OA) as depicted in Figure 1A and 1B, respectively (Gould 1971). These lateral transposition patterns result from dissociations between growth and development in the ontogeny of descendants (Gould 1971). Geometric scaling may be necessary for phyletic size changes when traits exhibit strong OA otherwise scaling up or down an ancestral ontogeny (i.e., “ontogenetic scaling”) may produce maladaptive trait forms (Kurtén 1954, 1955; Gould 1966, 1971). However, a disadvantage of geometric scaling is that it may produce trait sizes that have reduced biomechanical performance (Gould 1971).

**Figure 1.**
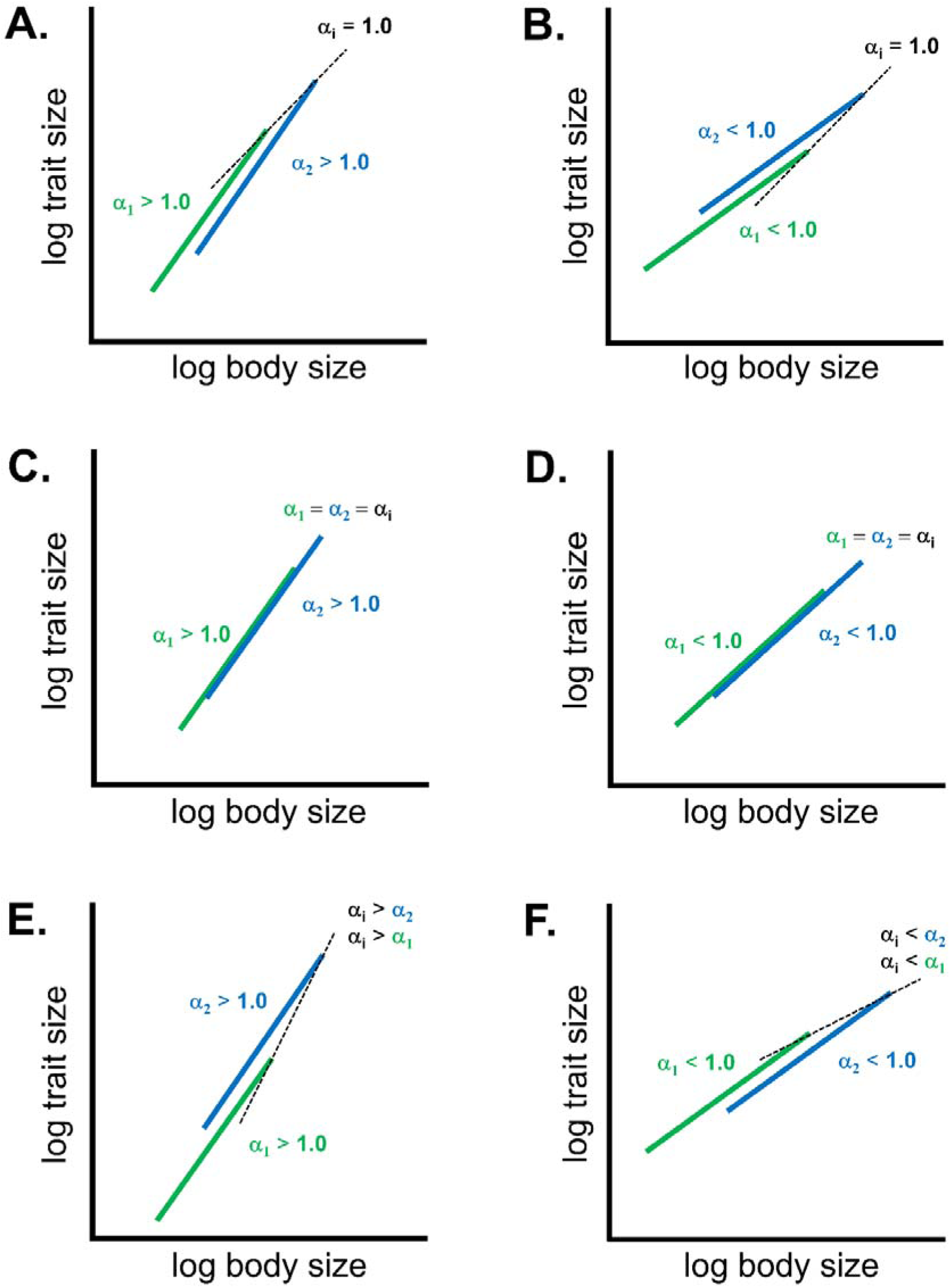
Major trait scaling patterns in pairwise comparisons of ontogenetic (or static) allometric trajectories. (A) Dissociation between ancestral and descendant trajectories leading to geometric similarity (GS) of positive ontogenetic or static allometric (OA) traits. This occurs when both trajectories’ slopes exhibit positive OA (i.e., α_1_ > 1 and α_2_ > 1) and the interpopulational or interspecific slope (α_i_) reflects GS (i.e., α_i_ = 1). (B) Dissociation between ancestral and descendant trajectories leading to geometric similarity (GS) of negative OA traits. This occurs when both trajectories’ slopes display negative OA (i.e., α_1_ < 1 and α_2_ < 1) and α_i_ = 1. (C) Truncation or extension of ancestral trajectory leading to positive evolutionary allometry (EA) via ontogenetic scaling (i.e., α_i_ > 1). This occurs when a descendant trajectory scales down or up an ancestral trajectory resulting in partially overlapping trajectories. (D) Truncation or extension of ancestral trajectory leading to negative EA via ontogenetic scaling (i.e., α_i_ < 1). This occurs when a descendant trajectory scales down or up an ancestral trajectory resulting in partially overlapping trajectories. (E) Transposition of ancestral and descendant trajectories leading to positive EA of positive OA traits (i.e., α_i_ > 1). This occurs when both trajectories’ slopes display positive OA (i.e., α_1_ > 1, α_2_ > 1) and α_i_ is greater than α_1_ and α_2_. (F) Negative EA via transposition. This occurs when both trajectories’ slopes display negative OA (i.e., α_1_ < 1, α_2_ < 1) and α_i_ is less than α_1_ and α_2_. Blue and green lines represent trajectories of log trait size against log body size for two conspecific populations or closely related species. One trajectory in each comparison is the same as the ancestral trajectory and isometry is assumed to equal one. Dotted lines depict interpopulational or interspecific allometries between the largest adults in each population or species.

Ontogenetic scaling, which is a form of evolutionary allometry (hereafter EA), occurs when an allometric trait scales up or down an ancestral ontogeny (Gould 1971; Klingenberg 1998). This scaling pattern reflects positive or negative EA depending on whether the trait displays positive or negative OA (Figure 1C and 1D, respectively). Positive and negative EA can also arise when two trajectories are transposed, though the pattern depends on a trait’s OA (Figure 1E and 1F). Note that changes in slopes of trajectories can produce other spatial patterns on log-log plots. However, such changes in the direction of trajectories usually exhibit more subtle spatial patterns compared to the patterns depicted in Figure 1 (Klingenberg 1998).

Szarski (1964, pp. 124-125) hypothesized that divergence in allometric traits begins with geometric scaling followed by evolutionary allometric scaling as Gould (1971, p. 132) had noted. Implicit in this hypothesis is that directional natural selection drives evolutionary allometric scaling during the second (trait “re-optimization”) step of the process. Using amphibian metabolism to illustrate this idea, Szarski hypothesized that if one species increased in body size while a closely related species remained the size of their most recent common ancestor, then the following process would occur. First, the surface area of respiratory organs would undergo geometric scaling in the former species and thus the interspecific regression slope should attain an isometric value of 0.66 for the metabolism-body size scaling relationship. Second, the trait would then go through a positive EA stage, which, according to Szarski, would lead to compensatory changes in its shape by increasing its surface area of respiratory organs. Despite its potential broad importance to evolutionary biology, this trait divergence hypothesis has not received further evaluation since its inception over six decades ago.

More recently, the present author, unaware of Szarski’s hypothesis, articulated a somewhat similar trait divergence hypothesis based on new evidence. The preliminary findings, in part, suggested that growth and development became uncoupled in the ontogenies of several species of pygopodid lizards that underwent body size changes (see Supplementary File 1), which fits the hypothesis of geometric scaling (Gould 1971). These findings complemented earlier results showing that positive EA scaling patterns characterized older interspecific divergences in the group (Jennings 2002). This trait divergence pattern has gone unnoticed in Darwin’s finches as well. Notice that in Figure 3 of Boag (1984) the two populations of *Geospiza fortis* show spatial patterns consistent with GS (Figure 1A) for all three beak traits (versus body weight) while more divergent species pairs mainly showed positive EA (Figure 1E). Knowing that the tails of pygopodid lizards and beaks of the finches exhibited positive OA according to Jennings (2002) and Boag (1984), respectively, the following novel hypothesis can explain this trait divergence pattern. First, these traits underwent geometric scaling with initial changes in body size followed by positive evolutionary allometric scaling. Specifically, functional analysis would suggest that directional natural selection increased trait sizes during the EA step when it was body size that initially increased, or selection decreased trait sizes when body size had earlier decreased. Although this new hypothesis superficially resembles Szarski’s hypothesis, they are not comparable because the former hypothesis describes how positive OA traits undergo initial divergence while the latter attempted to explain divergence of a negative OA trait (see Table 3 in Szarski 1964).

If the new hypothesis for divergence of positive OA traits is correct, then, contrary to Szarski’s hypothesis, negative OA traits should first undergo geometric scaling followed by *negative* evolutionary allometric scaling during the second step. This process is expected because, if directional selection re-optimizes negative OA traits during step 2, these traits should decrease in size when body size increased during step 1 or increase in size when body size originally decreased. In this paper, we develop a new model of allometric trait divergence based on the present author’s hypothesis. We then test five of the model’s key predictions using a dataset of locomotor-relevant trait data for pygopodid lizards. The results of this study support all predictions for positive and negative OA traits and thus contradict step 2 in Szarski’s hypothesis. The results also provide additional insights into trait divergence in pygopodids and other well-studied species.

## MATERIALS AND METHODS

### A scaling model of trait evolution

A model of the allometric trait divergence process referred to as the “geometric-similarity-first model” (GSF model) is developed here. Initial trait divergence depends on its OA (positive or negative) and the direction of a population’s body size change (increasing or decreasing) compared to its ancestral population. Thus, four fundamentally different evolutionary scenarios characterize the GSF model, each of which is outlined below.

When an ancestral population yields two descendant populations, one of which (i.e., population B) evolves a *larger* body size, the model predicts that there will be no change in trait shape in population B (Phase 1, Figure 2A). The hypothesis that trait divergence begins between populations rather than between species is justified by many examples of inter-populational GS in the literature (e.g., Gould 1971) and the preliminary evidence from pygopodid lizards and Darwin’s finches discussed in the introduction. This GS phase may continue beyond the time at which populations A and B evolve into species A and B, respectively, as has been observed in many studies (e.g., Gould 1966, 1971; Phase 2, Figure 2A). However, because species B presumably experienced decreased trait performance as a result of geometric scaling, directional natural selection may favor a compensatory *increase* in the trait’s size. The model predicts that the trajectory of species B should eventually overlap part of the trajectory of species A and thus show a “pseudo-ontogenetic scaling” pattern (Phase 3, Figure 2A). As observed above, however, the predominant scaling pattern among Darwin’s finches was positive EA via transposition, which suggests that pseudo-ontogenetic scaling may be a transitional form of EA. Thus, the strength of the EA relationship may increase as the trajectory of species B shifts further *upwards* until the trait attains an optimal shape (Phase 4, Figure 2A). Thus, positive EA characterizes the third and fourth phases of trait divergence.

**Figure 2.**
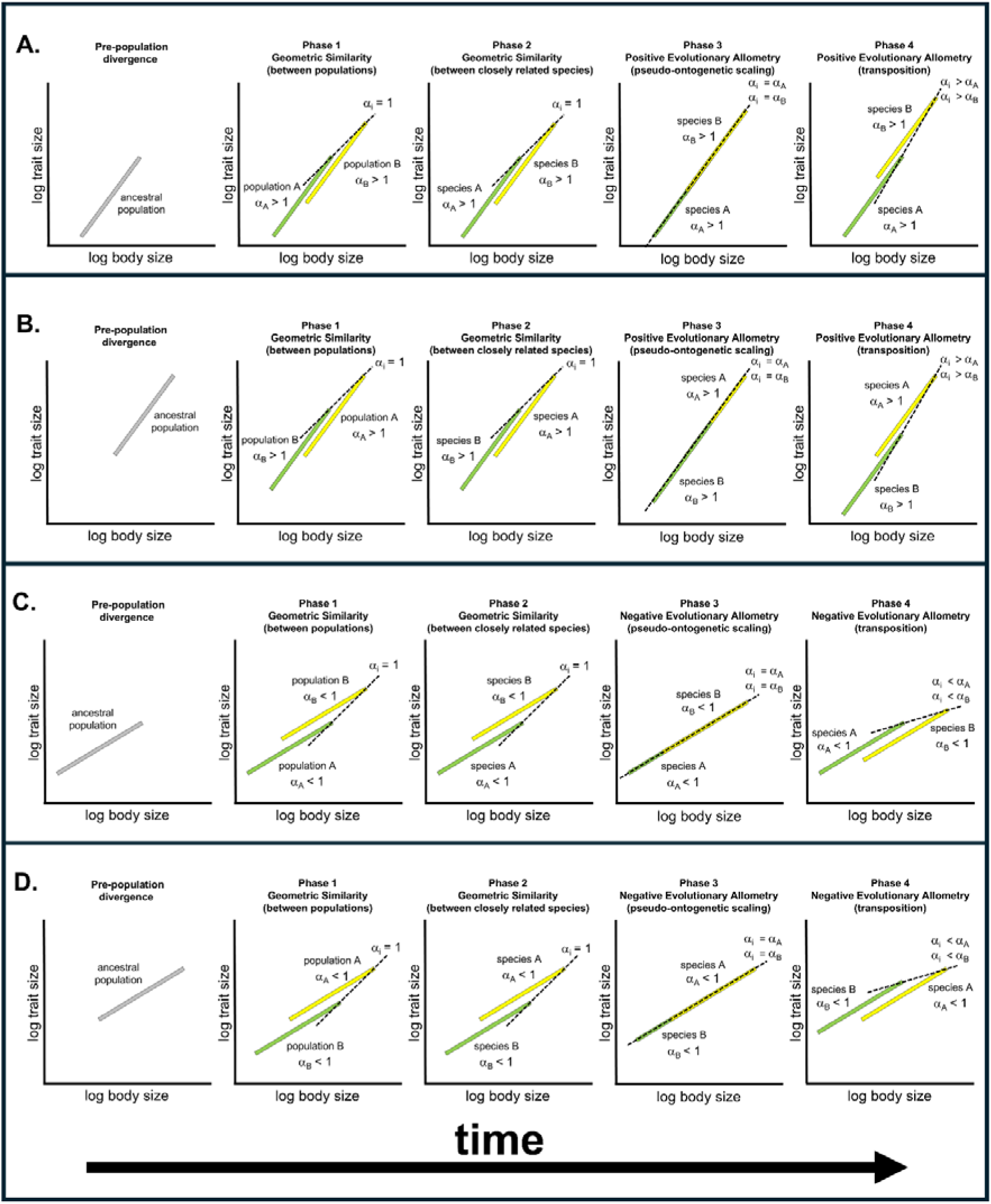
Geometric-similarity-first model of trait divergence. Each graph shows ontogenetic allometric trajectories plotted as log trait size on log body size. (A) Scenario 1: body size increases in population B (relative to the ancestor) and the trait shows positive ontogenetic allometry (OA). (B) Scenario 2: body size decreases in population B and the trait displays positive OA. (C) Scenario 3: body size increases in population B and the trait exhibits negative OA. (D) Scenario 4: body size decreases in population B and the trait shows negative OA. In each comparison the yellow trajectory has a larger maximum body size compared to the green trajectory. The population A and species A trajectories always resemble their ancestral trajectory (gray), whereas the population B and species B trajectories diverge from their ancestral trajectory. Intraspecific slopes (α_A_, α_B_) > 1 indicate positive OA while slopes < 1 signify negative OA. Dashed lines show interpopulational or interspecific scaling relationships (α_i_). Values of α_i_ equal to 1 indicate GS, α_i_ > 1 exhibits positive EA, and α_i_ < 1 means negative EA.

**Figure 3.**
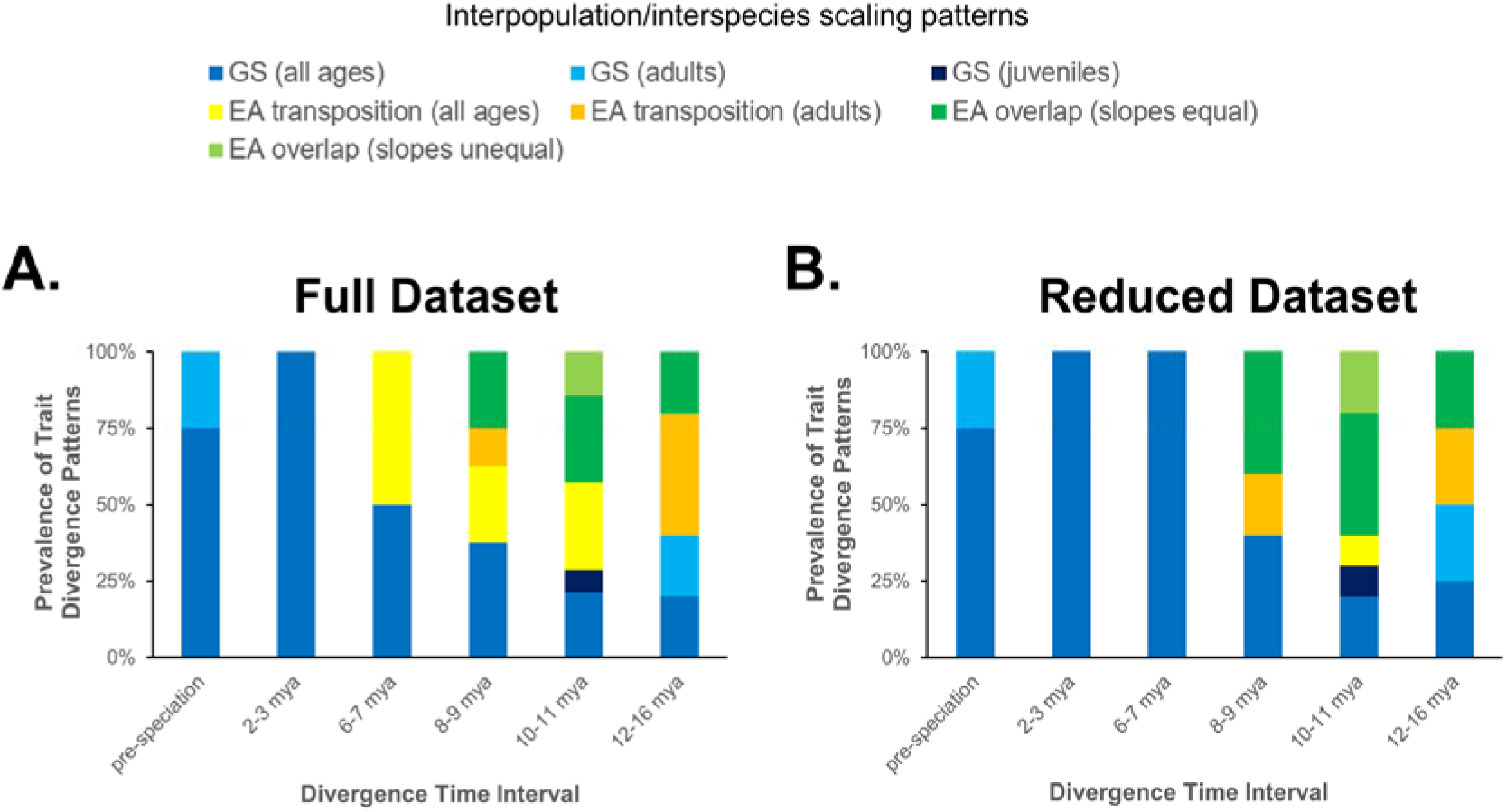
Test of the prediction that geometric similarity (GS) precedes evolutionary allometry (EA) during initial trait divergence using ln*Tail* on ln*SVL* pairwise comparisons. (A) Prevalence of trait divergence patterns during each divergence time interval using the full dataset. (B) Prevalence of trait divergence patterns during each time interval using the reduced dataset. Note the full dataset consisted of all OTU pairs that had one OTU with a longer maximum *SVL* compared to the other OTU, whereas the reduced dataset consisted only of OTU pairs in which one of the OTUs was larger than the other OTU in both *SVL* and *BW̄* body size measures. Mya = millions of years ago. “Pre-speciation” refers to the initial divergence time between intraspecific populations, which is assumed to have occurred more recently than the minimum speciation time for any species in this study (i.e., 2 Mya). See Supplementary Methods for definitions of scaling patterns and Supplementary Data for source data.

When an ancestral population gives rise to two descendant populations, one of which (population B) evolves a *smaller* body size, the model predicts no change in trait shape in population B (Phase 1, Figure 2B). Phase 2 is similar to the one in Scenario 1 (Phase 2, Figure 2B). However, this scenario predicts that trait size will undergo an evolutionary size *decrease*, resulting in a pseudo-ontogenetic scaling pattern (Phase 3, Figure 2B). In Phase 4, directional natural selection causes the trajectory of species B to shift further *downwards* until trait size is optimized (Phase 4, Figure 2B). Like the first scenario, Phases 3 and 4 in this scenario show positive EA.

When an ancestral population gives rise to two descendant populations, one of which (population B) evolves a larger body size, this model predicts no change in trait shape in population B (Phase 1, Figure 2C). Phase 2 is similar to Phase 2 in the previous scenarios (Phase 2, Figure 2C). However, unlike the previous two scenarios, Phases 3 and 4 in this scenario reflect negative EA. In Phase 3, this model predicts that trait size *decreases*, resulting in partially overlapping trajectories (Phase 3, Figure 2C). In Phase 4, the trajectory of the larger species shifts further *downwards* until the trait is optimized (Phase 4, Figure 2C).

Lastly, when an ancestral population gives rise to two descendant populations, one of which (population B) evolves into a smaller body size, this model predicts that there will be no change in trait shape in population B (Phase 1, Figure 2D). Phase 2 is similar to Phase 2 in the previous scenarios (Phase 2, Figure 2D). In Phase 3 of this scenario, the model predicts trait size will *increase* to achieve partially overlapping trajectories and thus display a pseudo-ontogenetic scaling pattern (Phase 3, Figure 2D). In Phase 4, the larger species’ trajectory shifts further *upwards* until trait size is optimized (Phase 4, Figure 2D). Thus, this scenario also shows negative EA.

At least five predictions are deduced from the GSF model: 1) initial trait divergence between conspecific populations is characterized by GS; 2) GS is a temporary phase of trait divergence, which begins prior to speciation and ends soon after speciation; 3) all allometric traits undergo geometric scaling at the outset of trait divergence whenever body size changes in a population; 4) traits that exhibit positive OA undergo positive EA in the second step of the model (assuming predictions 1-3 are correct); and 5) traits that display negative OA undergo negative EA in the second step of the model (assuming predictions 1-3 are correct). These predictions can be tested using trait data for fossils, contemporary species, or a combination of the two. Paleontological data have the advantage that fossils are often dated, which means that investigators can determine the direction of body size change (e.g., Kurtén 1954) and therefore distinguish between trait divergence scenarios that only differ in the direction of body size change (e.g., between Figure 2A and B). For neontological data, comparative methods (Harvey and Pagel 1991) can be used to infer the body size of the most recent common ancestor for each pair of populations or closely related species in a pairwise comparison of allometric trajectories. Note that it is not necessary to know the direction of body size change to test these predictions. However, an estimate of the maximum body size of each population or species is required.

### Study organisms

The GSF model was tested using morphological data for pygopodid lizards, a group of at 51 species that are endemic to Australia and New Guinea (Jennings 2021; Pepper et al. 2025). Pygopodids are ideal for this purpose because all species groups are well-defined, the phylogenetic relationships are well resolved, and divergence times are available for most species (Jennings 2003; Brennan et al. 2016; Brennan and Oliver 2017; Skipwith et al. 2019; Jennings 2021; Pepper et al. 2025). Moreover, each species group represents an independent replicate of the initial trait divergence process. This group also contains lineages that show the hallmarks of adaptive radiation (e.g., sympatry of congeners, ecomorphological diversity; Schulter 2000). For instance, the genus *Delma*, which accounts for half the group’s species richness shows remarkable variation in relative tail lengths (i.e., 244-450% of their body lengths), which often correlates with habitat-substrate use (Kluge 1974; Storr et al. 1990; Cogger 1992; Ehmann 1992; Bamford 1998; Jennings et al. 2002, 2003; Maryan et al. 2007; Pianka 2010; Jennings unpublished data). In contrast, other lineages exhibit characteristics of a non-adaptive radiation (e.g., allopatric or parapatric distributions and ecomorphological uniformity; Gittenberger 1991). The 15 species of *Aprasia* fit this diversification profile as they show little variation in their morphologies and in their habitat use (all are fossorial in habit), and they only exhibit parapatric or allopatric distributions (Storr et al. 1990; Jennings et al. 2003; Maryan et al. 2013a, b; Maryan et al. 2015a). Pygopodids also show dramatic variation in interspecific body sizes (Jennings 2002; Garcia-Porta and Ord 2013).

### Trait data

This study is based on morphological data from 1,756 museum specimens representing 31 pygopodid species. All specimens were housed in the following natural history collections: Western Australian Museum, Queensland Museum, South Australian Museum, Australian Museum, Museums Victoria, Museum and Art Gallery of the Northern Territory, and the Museum of Comparative Zoology (Harvard University). Five linear measurements to the nearest millimeter (mm) using dial calipers were recorded from each specimen: snout-vent length (*SVL*), tail length (*Tail*), body width at the ear (*BW1*), body width in the middle of the trunk (*BW2*), and body width at the vent (*BW3*). These measurements were selected because pygopodids are effectively limbless, and thus body size and shape as well as tail length determines the locomotor performance of each species. Tails were only measured if they were original (i.e., non-regenerated). All morphometric data in this study are from Jennings (2025).

### Analyses

#### Definition of OTUs

The operational taxonomic units (OTUs) in this study usually corresponded to recognized species. However, multiple population-level OTUs for some species were identified using the following criteria: (1) if a species comprised multiple geographically disjunct populations or subspecies, then each one was defined as an OTU; and (2) if a species had a widespread distribution that spanned multiple biomes, then the sub-population for each biome was defined as a separate OTU. Accordingly, some species consisted of a population in tropical regions (i.e., “wet biome”) and a population in arid ecosystems (i.e., “dry biome”).

#### Testing the null hypothesis of isometry

Only traits demonstrating OA are suitable for analysis with the GSF model. Post-natal growth trajectories must therefore be estimated for all OTUs so that their slopes can be tested against the null hypothesis of isometry. Three trait scaling relationships were examined. The first was ln*Tail* regressed on ln*SVL*, which describes post-natal changes in relative tail length using *SVL* as the body size reference. The second was ln*Tail* on lnB^---^W^--^, which is relative tail length using B^---^W^--^, the average body width (*BW1* + *BW2* + *BW3*)/3, as the size reference. The third was lnB^---^W^--^ on ln*SVL*, which is relative body girth using *SVL* as the size measure. Growth trajectories for OTUs were estimated using linear regression (see Supplementary Methods).

#### Pairwise comparisons of post-natal growth trajectories

Because a major focus of this study was to examine initial trait divergence, pairwise comparisons of trajectories were limited to conspecific populations and species belonging to the same species group (see Jennings 2021 and Pepper et al. 2025 for more details about these groups). To determine whether a pair of trajectories partially overlapped with each other (i.e., exhibited ontogenetic or pseudo-ontogenetic scaling) or were transposed (i.e., showed GS or EA via transposition) it was necessary to test the null hypotheses of equal slopes and equal elevations using standard statistical methods (see Supplementary Methods). Eight scaling patterns exhibited by paired trajectories on ln-ln plots were defined: 1) GS (all ages), 2) GS (adults only), 3) GS (juveniles only), 4) EA overlap (equal slopes), 5) EA overlap (unequal slopes), 6) EA transposition (all ages), 7) EA transposition (adults only), and 8) EA transposition (juveniles only). See Supplementary Methods for definitions of these patterns.

Although *SVL* has long been the body size reference for lizards and snakes, an unforeseen complication arose during the analyses of maximum body size in pygopodid OTUs. In some pairs the OTU with the smaller maximum *SVL* had a larger maximum B^---^W^--^than the OTU having the larger maximum *SVL*. Given the uncertainty as to which measure of body size may be more appropriate to use in these cases, two versions of each dataset were constructed for testing the model’s predictions. The “full datasets” each contained all OTU pairs that had one OTU with a larger maximum *SVL* compared to the other OTU in each comparison, while the “reduced datasets” contained OTU pairs that had one OTU with a larger maximum *SVL and* larger B^---^W^--^ compared to the other OTU in the comparison.

## RESULTS

### Testing the null hypothesis of ontogenetic isometry

The average slope for 32 OTUs in the ln*Tail* on ln*SVL* regressions was 1.28 (0.88 - 1.56; Table S1). Of these slopes, 25 (78%) reflect statistically significant positive OA. Average *R*^2^ is 0.89 (0.52 - 0.99). These results show a marked tendency towards positive OA in tail length when *SVL* is the body size measure. The average slope across 32 OTUs in the ln*Tail* on ln*BW̄*regressions is 1.44 (0.81 - 1.87; Table S2). Only two slopes are less than or equal to the null value of one. Of these slopes, 26 (81%) exhibit significant positive OA. Average *R*^2^ is 0.83 (0.57 - 0.98). These results show pronounced positive OA in tail length when average B^---^W^--^ is the size reference. The average slope across 35 OTUs in the lnB^---^W^--^ on ln*SVL* regressions is 0.76 (0.45 - 0.90; Table S3). All OTUs have slopes that reflect negative OA, of which 33 are statistically significant. Average *R*^2^ is 0.83 (0.57 - 0.95).

### Scaling relationships

Pairwise comparisons of OTUs within each species group yield 126 unambiguous scaling patterns when *SVL* is the body size measure (Tables S4-S10 and Figures S1-S7). A summary of these results is shown in Table 1.

**Table 1.** Frequencies of scaling patterns for three locomotor-relevant traits within each pygopodid lizard species group. Scaling patterns for ln*Tail* on ln*SVL*, ln*Tail* on ln*BW̄*, and ln*BW̄* on ln*SVL* were derived from plots of pairwise comparisons of ontogenetic trajectories. See Supplementary Tables S4-S10 for source data and Supplementary Methods for definitions of scaling patterns. GS = geometric similarity; EA = evolutionary allometry.

| Scaling pattern | Species Group |  |  |  |  |  |  |  |  |
| --- | --- | --- | --- | --- | --- | --- | --- | --- | --- |
|  | repens | SE Aprasia | australis | impar | nasuta | tincta | Lialis | lepidopodus | nigriceps |
| GS (all ages) | 8 | 2 | 0 | 0 | 11 | 4 | 3 | 3 | 8 |
| GS (adults) | 2 | 0 | 0 | 0 | 0 | 0 | 3 | 0 | 3 |
| GS (juveniles) | 0 | 0 | 0 | 0 | 1 | 0 | 0 | 0 | 0 |
| EA overlap (equal slopes) | 6 | 2 | 1 | 3 | 7 | 1 | 0 | 0 | 4 |
| EA overlap (unequal slopes) | 0 | 1 | 2 | 2 | 0 | 1 | 0 | 0 | 2 |
| EA transposition (all ages) | 2 | 10 | 0 | 3 | 9 | 0 | 2 | 0 | 11 |
| EA transposition (adults) | 0 | 2 | 0 | 1 | 2 | 0 | 1 | 0 | 0 |
| EA transposition (juveniles) | 0 | 1 | 0 | 0 | 0 | 0 | 0 | 0 | 2 |

Of these pairs, 48 exhibit GS-type patterns while 78 reflect EA-type patterns. Variation in scaling patterns is evident within species groups, which suggests that ontogenetic trajectories have been evolutionarily labile, even between conspecific populations.

### Testing the predictions of the GSF model

#### Prediction 1: GS precedes EA

Figure 3 shows the prevalence of scaling patterns for ln*Tail* on ln*SVL* relationships by divergence time interval. For the full dataset, 75% of the pre-speciation (interpopulational) pairs exhibit GS (all ages) while the other 25% show GS (adults), respectively (Figure 3A). OTU pairs within the next older divergence time interval (i.e., 2-3 Mya) only show GS (all ages). Results based on the reduced dataset mirror the full dataset results (Figure 3B). These findings suggest that GS preceded EA for ln*Tail* on ln*SVL* in pygopodid lizards.

Figure 4 shows the prevalence of scaling patterns for ln*Tail* on lnB^---^W^--^ relationships by divergence time interval. Results based on the full dataset (Figure 4A) show that 100% of the interpopulational pairs exhibit GS (all ages). Moreover, during the 2-3 Mya interval 2/3 of the OTU pairs also exhibit GS (all ages) while 1/3 show EA overlap (slopes equal). However, closer evaluation of these results reveals that two GS pairs (*Delma butleri*/*D. haroldi* and *D. pax*/*D. desmosa*) arose ∼2 Mya while the EA overlap pair (*Aprasia pseudopulchella*/*A. parapulchella*) underwent speciation earlier at ∼3 Mya. Thus, GS patterns prevailed from the start of trait divergence until ∼2 Mya. Results for the reduced dataset are similar to the full dataset results with respect to the prevalence of GS-type patterns (Figure 4B). This suggests that tail length in pygopodids initially changed with body width via geometric scaling.

**Figure 4.**
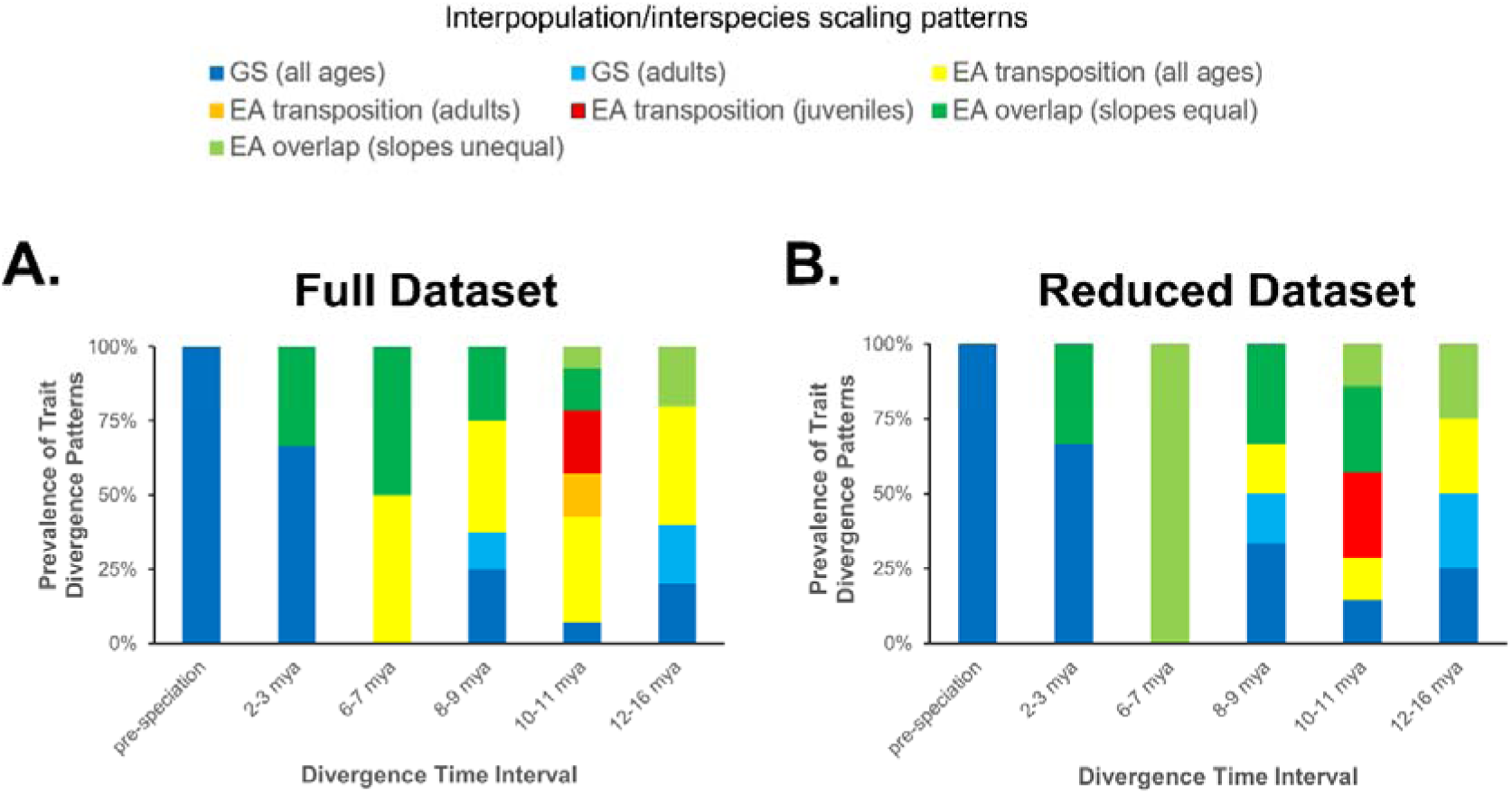
Test of the prediction that geometric similarity (GS) precedes evolutionary allometry (EA) during initial trait divergence using ln*Tail* on *BW̄* pairwise comparisons. (A) Prevalence of trait divergence patterns during each divergence time interval using the full dataset. (B) Prevalence of trait divergence patterns during each time interval using the reduced dataset. Note the full dataset consisted of all OTU pairs that had one OTU with a longer maximum *SVL* compared to the other OTU, whereas the reduced dataset consisted only of OTU pairs in which one of the OTUs was larger than the other OTU in both *SVL* and *BW̄* body size measures. Mya = millions of years ago. “Pre-speciation” refers to the initial divergence time between intraspecific populations, which is assumed to have occurred more recently than the minimum speciation time for any species in this study (i.e., 2 Mya). See Supplementary Methods for definitions of scaling patterns and Supplementary Data for source data.

Figure 5 shows the prevalence of scaling patterns for ln*BW̄* on ln*SVL* relationships by divergence time interval. Results for the full dataset show that 50% of the interpopulational pairs exhibit GS (all ages) while the other 50% display GS (adults only), respectively (Figure 5A). Moreover, during the 2-3 Mya interval 2/3 of the OTU pairs also display GS (all ages) while 1/3 show EA via transposition (all ages). However, divergence times suggest that the two GS pairs (*Delma butleri*/*D. haroldi* and *D. pax*/*D. desmosa*) arose ∼2 Mya while the EA transposition pair (*Aprasia pseudopulchella*/*A. parapulchella*) underwent speciation earlier at ∼3 Mya. Thus, only GS-type patterns are apparent for OTU pairs with pre-speciation to 2 Mya divergence times. Results for the reduced and full dataset resemble each other except the seven OTU pairs in the 2-3 Mya time interval show GS-type patterns (Figure 5B). These findings suggest that GS rather than EA characterizes initial divergences in body width.

**Figure 5.**
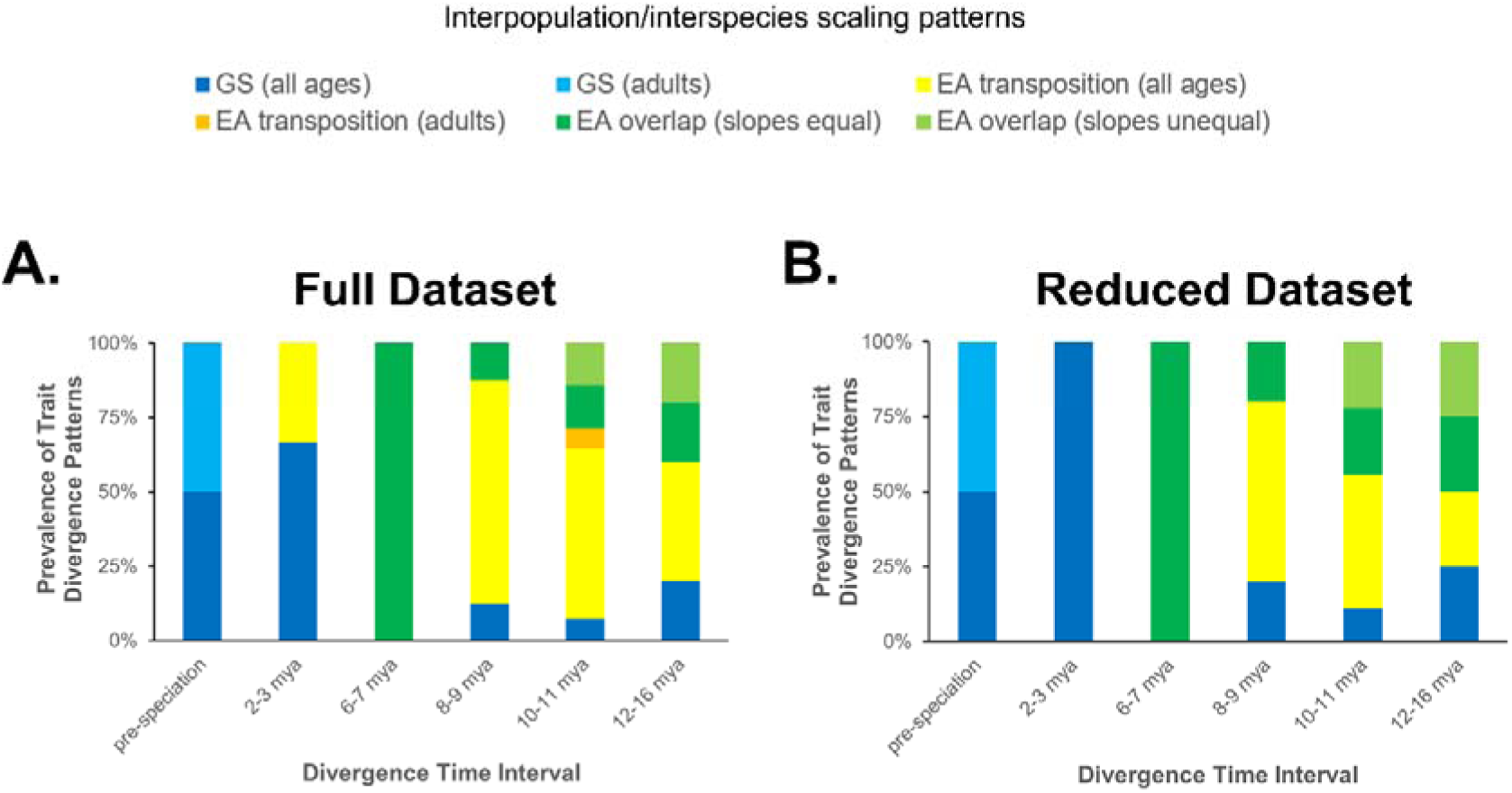
Test of the prediction that geometric similarity (GS) precedes evolutionary allometry (EA) during initial trait divergence using *BW̄* on ln*SVL* pairwise comparisons. (A) Prevalence of trait divergence patterns during each divergence time interval using the full dataset. (B) Prevalence of trait divergence patterns during each time interval using the reduced dataset. Note the full dataset consisted of all OTU pairs that had one OTU with a longer maximum *SVL* compared to the other OTU, whereas the reduced dataset consisted only of OTU pairs in which one of the OTUs was larger than the other OTU in both *SVL* and *BW̄* body size measures. Mya = millions of years ago. “Pre-speciation” refers to the initial divergence time between intraspecific populations, which is assumed to have occurred more recently than the minimum speciation time for any species in this study (i.e., 2 Mya). See Supplementary Methods for definitions of scaling patterns and Supplementary Data for source data.

*Prediction 2: Geometric similarity is a temporary phase of trait divergence* For all three locomotor-relevant traits, the prevalence of GS-type patterns decline as divergence times increase, whereas the prevalence of EA-like patterns increases as divergence times increase (Figures 3-5). These observations are consistent with the idea that GS is a temporary phase of allometric trait divergence rather than a stable evolutionary end point. Moreover, all interpopulational OTU pairs exhibit some form of GS for all three locomotor-relevant traits just before and after the time of speciation (Figures 3-5).

*Prediction 3: all allometric traits undergo geometric scaling with size changes* If we examine the ln*Tail* on ln*SVL*, ln*Tail* on ln*BW̄*, and ln*BW̄* on ln*SVL* scaling relationships for OTU pairs that have divergence time estimates, all four OTU pairs with pre-speciation-level divergences and two of the three pairs with 2-3 Mya divergence times only display GS-type patterns (Figure 6A). These results reveal that the two OTU pairs with ∼2 Mya divergence times (*Delma butleri*/*D. haroldi* and *D. pax*/*D. desmosa*) only show GS-type patterns while a third OTU pair that had a ∼3 Mya divergence time (*Aprasia pseudopulchella*/*A. parapulchella*) exhibits a mixture of GS and EA patterns. Therefore, all six OTU pairs with divergence times ranging from pre-speciation to ∼2 Mya only show GS-type patterns. However, another OTU pair, *L. burtonis* (wet biome) vs. *L. jicari*, shows GS-type patterns for all scaling relationships even though its estimated divergence time is 16 Mya—the oldest divergence time for any OTU pair in this study. Viewed another way, of the 30 pairs with divergence times greater than ∼2 Mya, 28 displayed either a mixture of GS and EA patterns or exclusively EA patterns. The reduced dataset results reveal that all OTU pairs in the pre-speciation and 2-3 Mya divergence time intervals only show GS-type patterns (Figure 6B). These results show that the initial stage of trait divergence is characterized by GS and thereby supports this prediction.

**Figure 6.**
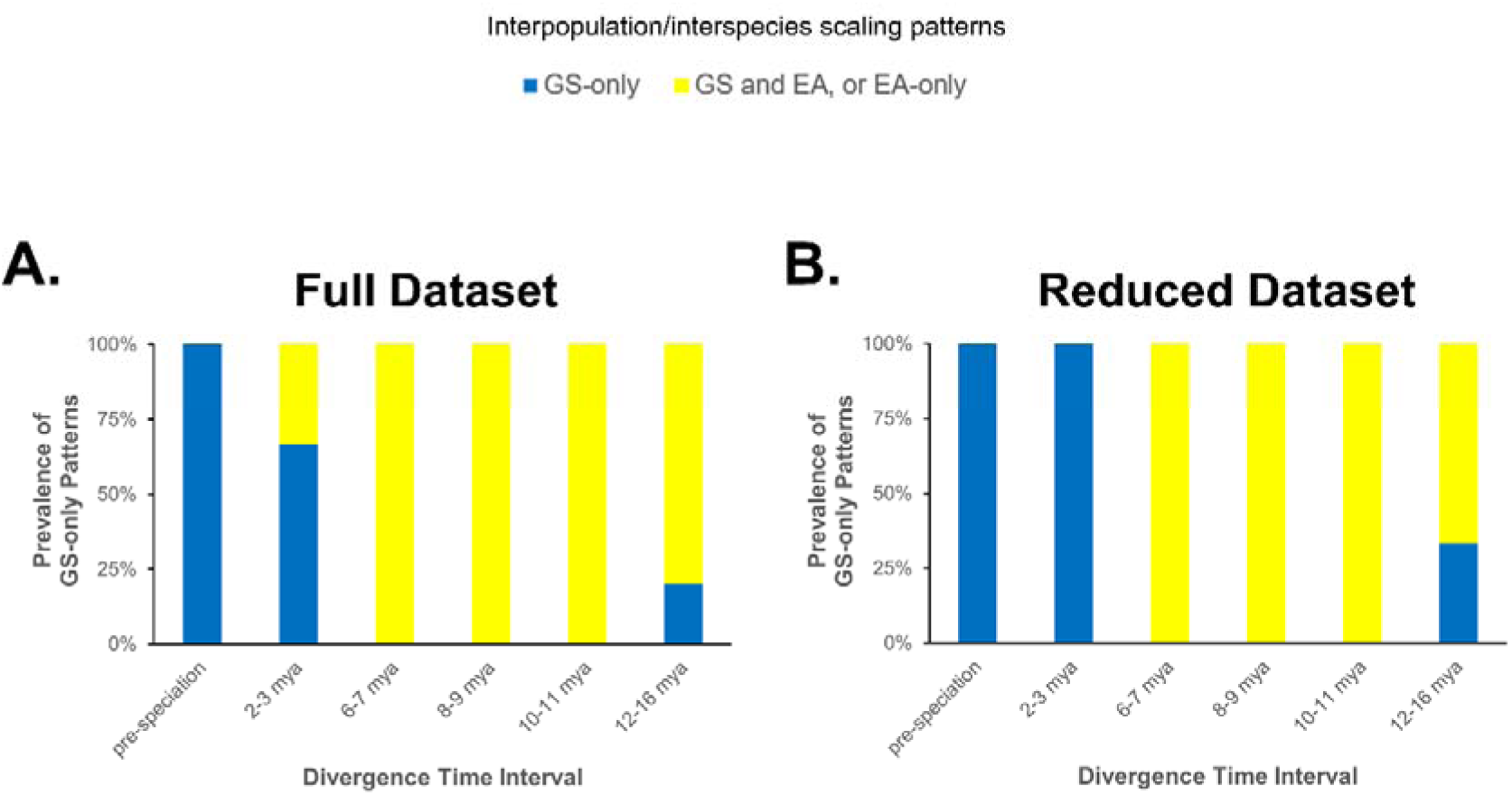
Test of the prediction that all allometric traits display geometric similarity (GS) during initial trait divergence. (A) Prevalence of GS-only scaling patterns (i.e., for ln*Tail* on ln*SVL*, ln*Tail* on ln*BW̄*, and ln*BW̄* on ln*SVL*) using the full dataset. Each bar shows the prevalence of OTU pairs that only showed GS patterns (blue) vs. OTU pairs that did not exclusively exhibit GS patterns (yellow). (B) Prevalence of GS-only scaling patterns (i.e., for ln*Tail* on ln*SVL*, ln*Tail* on ln*BW̄*, and ln*BW̄* on ln*SVL*) using the reduced dataset. Note the full dataset consisted of all OTU pairs that had one OTU with a longer maximum *SVL* compared to the other OTU, whereas the reduced dataset consisted only of OTU pairs in which one of the OTUs was larger than the other OTU in both *SVL* and *BW̄* body size measures. Mya = millions of years ago. “Pre-speciation” refers to the initial time of divergence between intraspecific populations, which is assumed to have occurred more recently than the minimum speciation time for any species in this study (i.e., 2 Mya). See Supplementary Data for source data.

*Prediction 4: Positive OA traits undergo positive EA following the GS phase* If we assume that the six pygopodid OTU pairs with ≤ 2 Mya divergence times exemplify an initial GS stage of trait evolution as the above results suggest, then predictions concerning the hypothetical second stage of trait divergence can be tested. Specifically, data for the 30 other longer diverged (i.e., > 2 Mya) OTU pairs that show unambiguous scaling patterns for the two positive OA traits are of use here (see Supplementary Data). Of these latter pairs, 19 (63%) exhibit positive EA for the ln*Tail* on ln*SVL* relationship. Similarly, 24/30 (80%) patterns for the ln*Tail* on ln*BW̄* relationship also reflect positive EA-type patterns. These observations are largely consistent with this model prediction.

Prediction 5: Negative OA traits undergo negative EA following the GS phase

The lnB^---^W^--^ on ln*SVL* relationship reflects negative OA. Accordingly, we can use this trait’s scaling patterns for the same 30 OTU pairs to test this prediction (see Supplementary Data). Of these pairs, 27 (90%) show negative EA patterns for this trait thereby supporting this prediction.

## DISCUSSION

### Empirical support for step 1 of the GSF model

Geometric scaling may be a developmental mechanism that permits organisms to evolve rapid body size changes while largely maintaining constant organ shapes (Gould 1971). Without such a mechanism, it is thought that evolutionary allometric scaling caused by increases or decreases in a population’s body size might produce maladaptive trait shapes (Kurtén 1954). While numerous instances of GS are documented in the literature, it is still uncertain whether GS represents a fundamental initial phase in allometric trait divergence. In the present study of locomotor-relevant traits in the snake-like pygopodid lizards, GS-type patterns were consistently observed for pairs of conspecific populations and the most-recently originated species in each of six independent species groups. This is compelling evidence that growth and development of allometric traits became dissociated between ancestral and descendant trajectories at the outset of trait divergence. Moreover, EA scaling patterns were only evident for longer-diverged species in each group. These results corroborate the first step of the GSF model, which posits that GS precedes EA during the process of trait divergence.

The pygopodid trait data also demonstrated that GS is a temporary phase of trait evolution as predicted by the model. Although these data showed that GS is a “temporary” phase of trait divergence, the duration of this phase—lasting more than several million years—suggests that traits with suboptimal sizes performed well enough to allow those organisms to survive and reproduce over the long term. Nonetheless, the GSF model predicts that GS will eventually transition into an EA phase as directional natural selection (e.g., intraspecific competition or predation) hones the shapes of these traits in the incremental manner first envisioned by Darwin.

What remains to be explained are the upticks in the prevalence of geometrically similar OTUs in the oldest (12-16 Mya) divergence time category, which are intriguing exceptions to the dominant GS-precedes-EA pattern. One of these pairs*, Lialis burtonis* (wet biome population) and *L. jicari* exhibited more drastic morphological divergence than was observed for younger OTU pairs that displayed GS. Specifically, the trunks of *L. jicari* were tapered from mid-body towards the neck in a more extreme fashion than was observed in any specimens of *L. burtonis*, while the heads and snouts of the former species have become far narrower and elongated than was observed in the latter species. When one species undergoes such a reorganization of proportions its trajectory has become independent of the ancestral trajectory, a phenomenon termed “allometric escape” (Tonini et al. 2020). The resulting novel trait shapes may evolve strikingly new functions that enable species to occupy completely different niches than their closest relatives (Gould 1971; Grant 1986; Tonini et al. 2020). However, this type of trait evolution appears to only occur between highly divergent lineages (Gould 1971; Tonini et al. 2020) and thus likely does not occur during initial trait divergence.

Another prediction of the model holds that all traits showing OA must undergo geometric scaling with initial changes in body size. Except for the *L. burtonis* (wet biome) and *L. jicari* OTU pair, which apparently showed false positive GS patterns for all three traits, this prediction was upheld by the six pairs having the most recent divergence times. Moreover, the two tail length traits displayed the expected GS pattern (see Figure 1A) as did the body width trait (see Figure 1B).

How well does the GSF model fit the beak traits of Darwin’s finches? As Boag suggested (1984, p. 427), there may have been additional biologically meaningful transpositions hidden in Figure 3 of that paper that were not statistically supported. Accordingly, it may be fruitful to conduct visual comparisons of the different ground finch populations and species in that figure involving the three beak traits against body weight. Notice that 22/30 possible pairs show positive EA-like patterns while 8/30 pairs exhibit GS-like patterns (Table S11). While the three GS patterns for the two *G. fortis* populations fit the model as was noted in the introduction, the other five apparent cases of GS require explanation.

Four of the five species pairs that exhibited GS-like patterns included *G. scandens*, the cactus finch (see Table S11). This observation fits the larger pattern in Boag’s results, which showed that 12 out of the 20 statistically significant transpositions (see Table VIII in Boag 1984) involved the cactus finch. The morphological distinctiveness of this species is demonstrated by its specialized beak morphology, its adaptation to a cactus-based diet rather than the granivorous diets of other ground finches, and its deep phylogenetic divergence compared to the other ground finches in Boag’s study (Grant 1986; Grant and Grant 2008). Thus, like the pygopodid *L. jicari* the cactus finch beaks may represent a case of allometric escape rather than GS via initial trait divergence. Further support for this idea comes from a gene expression study, which showed that the gene for beak length was expressed at higher levels in the tips of cactus finch beaks—giving rise to their long and pointed beaks—compared to other ground finch species at the same stage of development (Abzhanov et al. 2006; Grant and Grant 2008). This leads us to conclude that the pair of *G. fortis* populations and the *G. magnirostris*/*G. fortis* pair represent the only manifestations of GS via *initial* divergence involving the beak traits thereby corroborating step 1 of the GSF model.

### Empirical support for step 2 of the GSF model

If traits do suffer a performance loss as a result of geometric scaling, then they may need to be restored to their functionally optimal shapes by directional natural selection. If this is correct, then it implies that following geometric scaling positive OA traits will undergo positive evolutionary allometric scaling while negative OA traits must go through negative evolutionary allometric scaling (see Figure 2). But is this what actually occurs in nature?

The pygopodid trait data provide support for these two predictions. While all conspecific populations and species formed less than ∼2 Mya only exhibited GS-type patterns in the two positive OA traits (i.e., relative tail length traits), species that originated more than ∼2 Mya tended to show positive EA in these traits. Moreover, the negative OA trait (relative body girth) exclusively manifested GS patterns in the recently originated (< 2 Mya) OTUs but displayed negative EA in 90% of the longer-diverged OTU pairs, which contradicts step 2 in Szarski’s (1964) trait divergence hypothesis.

How can we explain the pygopodid OTU pairs that did not agree with these two predictions? At least two explanations can account for these observations. First, some of these “aberrant” scaling patterns may reflect GS rather than the opposite form of EA predicted by the model. This might be expected for younger OTU pairs where GS may linger in some traits while other traits have already transitioned to the specific types (positive or negative) of evolutionary allometric scaling predicted by the model. As discussed earlier, allometric escape in the oldest OTU pairs can also explain some of these unexpected results.

### Choice of body size reference revisited

Although most pygopodid OTU pairs contained one member that exhibited both the longer maximum *SVL* and B^---^W^--^, size differentials in ten other pairs were ambiguous owing to one OTU having the longer maximum *SVL* while the other had the larger maximum B^---^W^--^.

However, several lines of evidence suggest that *SVL* is nonetheless the correct size reference for these lizards. First, there were negligible differences between the results obtained from the full and reduced datasets in all analyses. Second, the six most-recently diverged OTU pairs were not ambiguous in this regard, yet they provided much of the empirical support for step 1 of the model. Third, the basic conclusions would have remained the same had B^---^W^--^ been employed as the size reference instead of *SVL*. Note also that using B^---^W^--^ as the reference would have changed some EA via transposition patterns into GS patterns, which would have violated Gould’s (1971) GS rules for traits that show negative OA. Applying these rules here, a population that *increases* in *SVL* is expected to also *increase* in B^---^W^--^ to maintain GS while a population that *decreases* in *SVL* is predicted to *decrease* in B^---^W^--^ to maintain GS. Accordingly, two OTUs cannot be geometrically similar to each other if one of them has a larger *SVL* while the other has the larger B^---^W^--^. Lastly, the GSF model accounts for these ambiguous OTU pairs when *SVL* is used as the size reference. For instance, if a population undergoes a body size increase, then *SVL* and B^---^W^--^ for that population will initially *increase* during a GS phase while during the subsequent negative EA phase B^---^W^--^ is expected to *decrease* in size (Figure 2C). Alternatively, if the population undergoes a body size *decrease*, then *SVL* and B^---^W^--^ for that population will initially *decrease* during the GS phase while during the later negative EA phase B^---^W^--^ is expected to *increase* in size (Figure 2D).

### Ontogenetic scaling and the GSF model

Reported cases of ontogenetic scaling (e.g., Esquerré et al. 2017; Gray et al. 2019; Pavón-Vázquez et al. 2022) between closely related species pose a problem for the GSF model because the model only considers the hypothetical occurrence of “pseudo-ontogenetic scaling” and not true ontogenetic scaling. However, given the empirical support for the GSF model, we must ask the question: does true ontogenetic scaling exist? A corollary to Kurtén’s (1954) and Gould’s (1971) argument that traits showing “strong” OA must undergo GS with changes in body size is that traits showing “weak” OA are allowed to extrapolate up or down an ontogenetic trajectory to new body sizes without suffering adverse fitness consequences. However, at least two difficulties exist with this hypothesis. First, Lande (1985) argued that ontogenetic scaling cannot occur, at least when body size is the target of selection. Another complication is that ontogenetic scaling would require a developmental “switch” that determines whether trait size undergoes ontogenetic scaling or geometric scaling when a population changes in body size. If such a switch exists, then how strong does trait allometry need to be for one or the other scaling process to occur?

A simpler explanation can account for putative cases of ontogenetic scaling. Consider a scenario in which one population is ∼8% larger than another conspecific population. The 8% difference here corresponds to the minimum amount of body size difference observed between pygopodid populations that exhibited statistically supported GS patterns. In this scenario involving a trait showing positive OA, notice that ontogenetic scaling is difficult to distinguish from GS when OA is relatively weak, whereas a GS pattern becomes more and more distinctive as the strength of OA increases (Figure 7). In pygopodids, the average intraspecific *α* across all OTUs for the tail-scaling relationships ranged from 1.3 to 1.4, suggesting that tail length shows relatively strong OA. We can therefore reconcile the model with putative cases of ontogenetic scaling using two non-mutually exclusive arguments: 1) ontogenetic scaling and GS patterns may not be statistically distinguishable when traits exhibit weak OA; and 2) apparent cases of ontogenetic scaling may instead represent instances of pseudo-ontogenetic scaling, especially if studies did not include samples from different conspecific populations and nascent species.

**Figure 7.**
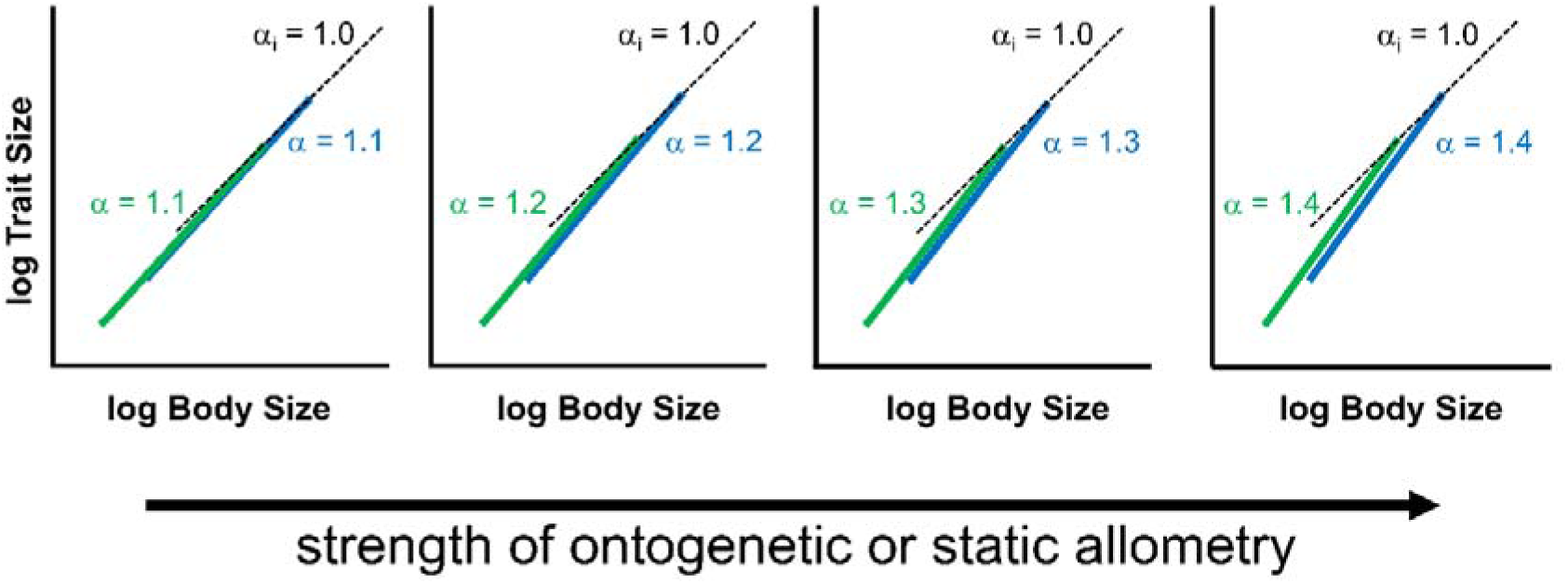
Strength of ontogenetic or static allometry (OA) and geometric similarity (GS) scaling patterns. Hypothetical log-log plots illustrate pairwise comparisons of ontogenetic (or static) trajectories for two populations (shown in green and blue) involving a trait that exhibits differing strengths of positive OA. The slopes, α, for the two populations are shown in green and blue, whereas the interpopulation slope, α_i_ (dashed line connecting the adults of both populations) is equal to 1.0 in all plots (i.e., GS). The trait shows differing strengths of positive OA ranging from α = 1.1 (weakest) to 1.4 (strongest) assuming isometry equals1.0. In each plot, the blue population underwent an 8% body size increase compared to the green population. This body size increase corresponds to the minimum body size difference for pygopodid populations that showed GS in locomotor traits.

### Implications

*Adaptive peak shifts through geometric scaling of viability traits* Locomotory adaptations are but one of several types of “viability traits” that enable organisms to exploit trophic and habitat niches located on an adaptive landscape (Bonduriansky 2011). How then did pygopodids undertake adaptive peak shifts that allowed them to occupy different niches? It is thought that closely related species must first attain some amount of “initial differences” while they are in allopatry so that ecological character displacement (Brown and Wilson 1956) can later cause further adaptive divergence between the two species once they are in sympatry, and hence maintain their coexistence (Lack 1947; Schluter 2000). The results of this study suggest that geometric scaling may be a general mechanism that produces these initial differences, but can this form of trait scaling facilitate a peak shift? The Delma tincta group may be a good model for understanding the evolution of initial trait divergences in adaptive radiation because this group contains pairs of species that exhibit allopatric and sympatric distributions. Similar to other recently diverged populations and sister species (i.e., those that diverged < 2 Mya), the D. tincta group pair *D. desmosa* and *D. pax* displayed an allopatric distribution and showed GS patterns for all three locomotor traits. In contrast, while group members *D. branchia* and *D. pax* show patterns that are consistent with GS for the three traits (see Supplementary File 2), these species are sympatric across the Pilbara region of Western Australia and are syntopic at many localities (Storr et al. 1990; Maryan et al. 2007; Pepper et al. 2025). Two other species in this group, *D. borea* and *D. desmosa*, show a mixture of scaling patterns, one of which is GS (see Table S8), and both species have been found at the same locality (Maryan et al. 2007). Interestingly, two sympatric species of Darwin’s finches, *G. fortis* and *G. magnirostris*, also display an instance of GS for one of the beak traits (see Table S11). These observations are noteworthy because they suggest that a sufficient amount of trait divergence occurred during initial divergence via geometric scaling when populations were in allopatry to later allow for the coexistence of closely related species. If this is correct, then character displacement—if it occurred—has been relegated to a finishing role in these trait divergences as Schluter (2000) had suggested might occur.

If geometric scaling facilitated peak shifts in pygopodid lizards, then what drove this initial trait divergence process? Genetic drift can move the mean population phenotype from one peak to another peak (Wright 1931, 1932) provided that adaptive valleys are shallow and population sizes are small (Schluter 2000). This hypothesis is unsatisfactory because most species of pygopodids exhibited expansive geographical distributions on the Australian continent (see Cogger 1992 and Ehmann 1992). This leaves three natural selection-based hypotheses to explain peak shifts in pygopodids: direct natural selection on Y (locomotory traits) may have caused correlated changes in X (body size), direct selection on X induced correlated responses in Y, and a combination of these two mechanisms (Gould 1966; 1974; Lande 1979; 1985; Price et al. 1993; Schluter 2000). The repeated independent evolution of wet and dry biome ecotypes in some pygopodid species is evidence that divergent natural selection caused trait divergence (Gould 1966; Schulter 2000), though this does not pinpoint the target(s) of selection.

It is counterintuitive that natural selection would drive geometric scaling by acting directly on a specific organ (e.g., beak)—and hence induce a change in body size as a correlated response—because the trait in question would initially evolve a suboptimal size relative to body size according to biomechanical analysis. However, if a new population is founded on an adaptive landscape that contains a different niche than was found on the source population’s landscape, then individuals in the population having smaller or larger trait sizes compared to the ancestral trait size may be better able to exploit the available niche compared to individuals having the ancestral trait size. If this is correct, then the *absolute* size of a trait may be adequate in the short-term to allow a population or species to gain a foothold in a new niche, while over subsequent evolutionary time directional selection (via intra- and interspecific competition) can re-optimize trait shape according to the GSF model. This may explain why some pygopodid populations and species have persisted for millions of years despite having locomotor traits of apparently suboptimal sizes relative to their respective body sizes. This may have also been the case with Darwin’s finches, as Schluter (2000) pointed out that adaptive landscapes for seed sizes and hardness differ among islands in the Galápagos archipelago. Thus, a population that colonizes an island with larger and harder seeds may evolve larger beak and body sizes or vice versa (Grant 1986).

Natural selection could have also driven peak shifts by targeting body size and hence inducing changes in the sizes (but not shapes) of pygopodid locomotor traits as correlated responses. Simulations in Price et al. (1993) suggested that peak shifts can readily occur via correlated response when either natural selection on X is strong or when the fitness valley is shallow. The finding that pygopodids can survive and reproduce with presumably suboptimal locomotor traits over long evolutionary time periods suggests that stabilizing selection on these traits was not particularly strong. If this is correct, then moderate to strong selection on body size may have moved population phenotypes across shallow fitness valleys to new peaks. Body size selection may be the primary driver of geometric scaling and hence peak shifts via correlated responses for at least two reasons. First, body size may be a frequent target of natural selection because there are many advantages to evolving larger or smaller body sizes (e.g., Rensch 1948; Szarski 1964; Gould 1966, Lande 1979, 1985; Hanken and Wake 1993). Moreover, such advantages may offset the disadvantages of evolving traits with reduced performance capabilities owing to geometric scaling (Gould 1966). Second, body size, rather than individual organ size, is more likely to be a target of natural selection simply because there are many more genes influencing body size than any one organ.

*Adaptive peak shifts through geometric scaling of sexually selected traits* The possibilities for peak shifts via correlated response multiply when we also consider sexually selected traits (Bonduriansky 2011). This is because sexually selected traits often exhibit static allometry for the adults (Bonduriansky and Day 2003; Bonduriansky 2007; Eberhard et al. 2018; Stroud et al. 2023) and therefore these traits are also expected to follow the GSF model. In one scenario, sexual selection on body size may cause peak shifts involving viability traits (Bonduriansky 2011). Although the viability traits in this scenario would presumably be of suboptimal size owing to geometric scaling, the preliminary results of this study suggest that a such a peak shift may still occur. Another scenario in which sexual selection may cause a peak shift involves direct selection on a sexual trait. For example, the dewlaps of adult male *Anolis* lizards, which tend to display positive static allometry, may function to signal the size of the lizard to reproductive females and rivals (Stroud et al. 2023). Sexual selection for males with larger dewlaps would favor the largest males in the population even though their dewlaps would be geometrically similar to those of smaller males (see Phase 1 in Figure 2A). This could have two important consequences: 1) this will likely lead to an evolutionary increase in body size, which can help drive speciation (Bonduriansky 2011); and 2) this will likely lead to changes in the absolute sizes of viability traits via correlated response and thus could cause a peak shift. Analogous peak shift scenarios could play out with genital characters in Poeciliid fishes, which show negative static allometry (Bonduriansky 2007). The interplay between body size and organs important to sexual selection and viability may also explain the diversification of birds of paradise (Frith and Beehler 1998).

The extinct Irish Elk (*Megaloceros giganteus*) has long interested evolutionary biologists because of its immense body size and enormous antlers. Despite previously voicing concerns that ontogenetic scaling can lead to maladaptive traits, Gould (1974) suggested that the Irish Elk had undergone this mode of trait evolution, driven by selection on body and antler sizes along a positive static allometric trajectory. However, Tsuboi et al. (2024) rejected Gould’s hypothesis because they found that ontogenetic scaling didn’t account for its relative antler size, which was larger than predicted by ontogenetic scaling. The GSF model can account for these new findings: when this species initially underwent body size increase, its antler size was smaller than static allometry would have predicted (see phase 1 in Figure 2A). But subsequent directional selection (e.g., due to intersexual selection; Gould 1974) could have caused an increase in antler size until it was transposed above the intraspecific allometric line (see phase 4 in Figure 2A).

*“Niche scaling”: a possible mechanism for peak shifts via geometric scaling* The results of this study showed that divergence of allometric traits follows a predictable evolutionary pathway: traits initially undergo geometric scaling and then they undergo evolutionary allometric scaling during a “re-optimization” phase of divergence. The results also revealed that trait divergence between allopatric or parapatric populations due to geometric scaling may be significant enough to facilitate their coexistence in the same community as separate species at a later time. If this is correct, then trait divergence solely by geometric scaling may be sufficient to allow a population or species to invade a new peak on an adaptive landscape. Given the ubiquity of population-level body size evolution in nature, geometric scaling may be a common mechanism of peak shifts and therefore play a major role in macroevolution as was first suggested by Gould (1971). However, adaptive radiation theory (Simpson 1944; Lack 1947; Simpson 1953; Grant 1986; Schluter 2000; Grant and Grant 2008, 2024; Losos 2009; Bonduriansky 2011; Gillespie et al. 2020) has overlooked the potential role that geometric scaling may play in peak shifts. Given the importance of this possible mechanism of peak shifts, we propose naming it “niche scaling.”

## CONCLUSION

The geometric-similarity-first model (or GSF model) aims to explain initial divergence and subsequent evolution of allometric traits in populations and closely related species. In step 1 of the model, traits undergo geometric scaling with a population-level decrease or increase in body size. In step 2, directional natural selection “re-optimizes” traits via one of two alternative pathways: 1) if the trait displays positive ontogenetic (or static) allometry (OA), then it is expected to undergo positive evolutionary allometric scaling; or 2) if the trait shows negative OA, then it is predicted to undergo negative evolutionary allometric scaling. A morphological data set for three locomotor-relevant traits in pygopodid lizards supported all five predictions of the GSF model and hence contradicts the second step in Szarski’s (1964) trait divergence hypothesis. The GSF model may clarify our understanding of trait divergence in other species. For instance, the model provides novel insights on beak divergence in Darwin’s finches and antler evolution in the Irish Elk.

This study produced two other findings of possible importance to macroevolution. First, the initial geometric similarity phase in pygopodid lizards lasted several million years, which suggests that organisms can survive and reproduce over long evolutionary time intervals despite having biomechanically deficient traits caused by geometric scaling. The second key finding was that closely related species of pygopodid lizard species were found to coexist together even though they exhibited geometric similarity in at least one of the examined viability traits. Notably, two species of Darwin’s finches also showed this pattern. These observations suggest that geometric scaling may have played a role in peak shifts in these two groups. It is hypothesized that natural selection can drive geometric scaling of viability and sexually selected traits and thus act as a mechanism for peak shifts via correlated response on an adaptive landscape (see Price et al. 1993 and Bonduriansky 2011), a process here named “niche scaling.” Given the universality of body size evolution in species, and the predictable consequences of these events according to the GSF model, niche scaling could be an important mechanism in adaptive radiation.

## Acknowledgements

This work began during the present author’s Ph.D. dissertation research but was only recently finished thanks to assistance from many institutions and individuals. Assistance was provided by the Department of Integrative Biology at the University of Texas at Austin, Museum of Comparative Zoology at Harvard University, and the Department of Evolution, Ecology & Organismal Biology at the University of California, Riverside. Access to specimens was provided by Andrew Amey and Patrick Couper (Queensland Museum), Paul Horner, Sue Horner, and Danielle Edwards (Museum and Art Gallery of the Northern Territory), Jane Melville and Katie Date (Museums Victoria), Ross Sadlier, Allen Greer, Dane Trembath, and Jodi Rowley (Australian Museum), Mark Hutchinson (South Australian Museum), Laurie Smith, Paul Doughty, and Kailah Thorn (Western Australian Museum), Jose Rosado, James Hanken, Stevie Kennedy-Golds, and Joe Martinez (Museum of Comparative Zoology, Harvard University [MCZ]). Funding for travel to museums in 1999 was provided by a Continuing Graduate Student Fellowship from the University of Texas at Austin while funding for trips between 2022 and 2023 to the MCZ and museums in Australia were provided by an Ernst Mayr Travel Grant from Harvard University and a Science, Discovery & Breakthrough Grant from Geeks without Frontiers (UCR-A01081-87924-44-NSWBJ). The author is also grateful for the assistance he received from his doctoral advisor, Eric R. Pianka, and dissertation committee members Robert Dudley, David Cannatella, Carl Gans, Theodore Garland, Jr., and David Hillis. Samuel Sweet, Michael and Margaret Potter, and Vivian Menezes Leandro provided much assistance and encouragement over the years, without which this study could not have been completed.

## DATA AVAILABILITY

The data underlying this article are available in the online supplementary material and in the Figshare repository under doi: 10.6084/m9.figshare.30479747.

