## Supplementary File 1 for "A scaling theory of trait evolution"

### Using ontogenetic allometric trajectories to identify interpopulational trait scaling patterns in pygopodid lizards

A preliminary analysis of interpopulational scaling patterns for tail length and snout to vent length in pygopodid lizards was performed using the pairwise ontogenetic allometric trajectory approach (e.g., Kurtén 1954). In the dataset of Jennings (2002), one species of pygopodid lizard, *Pygopus lepidopodus*, was represented by samples taken from two geographically well-separated populations that differed in maximum body size. Samples of *P. lepidopodus* from the central east coast of Australia (hereafter "Eastern Population") attained larger body sizes—measured as snout-vent length (*SVL*)—compared to the specimens collected from the central coast of Western Australia (hereafter "Western Population"; Supplementary File 1—Table 1). Post-natal allometric trajectories were estimated using linear regression of ln-transformed tail length (ln*Tail*) and ln-transformed snout-vent length (ln*SVL*) data separately for each population. The ln-transformed data points and regression lines were then plotted to show their relative positions in 2-D morphospace. The plot of the two *P. lepidopodus* populations showed overlapping trajectories during the juvenile growth stages but diverging trajectories during the adult growth stages of both populations (Supplementary File 1—Figure 1). Given that the tail lengths of the lizards in both populations exhibited positive ontogenetic allometry (i.e., both slopes were greater than the isometric value of 1.0)—and thus fit the strong trend in pygopodid lizards (Jennings 2002), the Y-shaped scaling pattern suggests that the adults of each population diverged from each other in a manner that is consistent with geometric similarity (i.e., dissociation between size and shape in the adult cohort). These results must be considered preliminary due to the low sample size in the Eastern population.

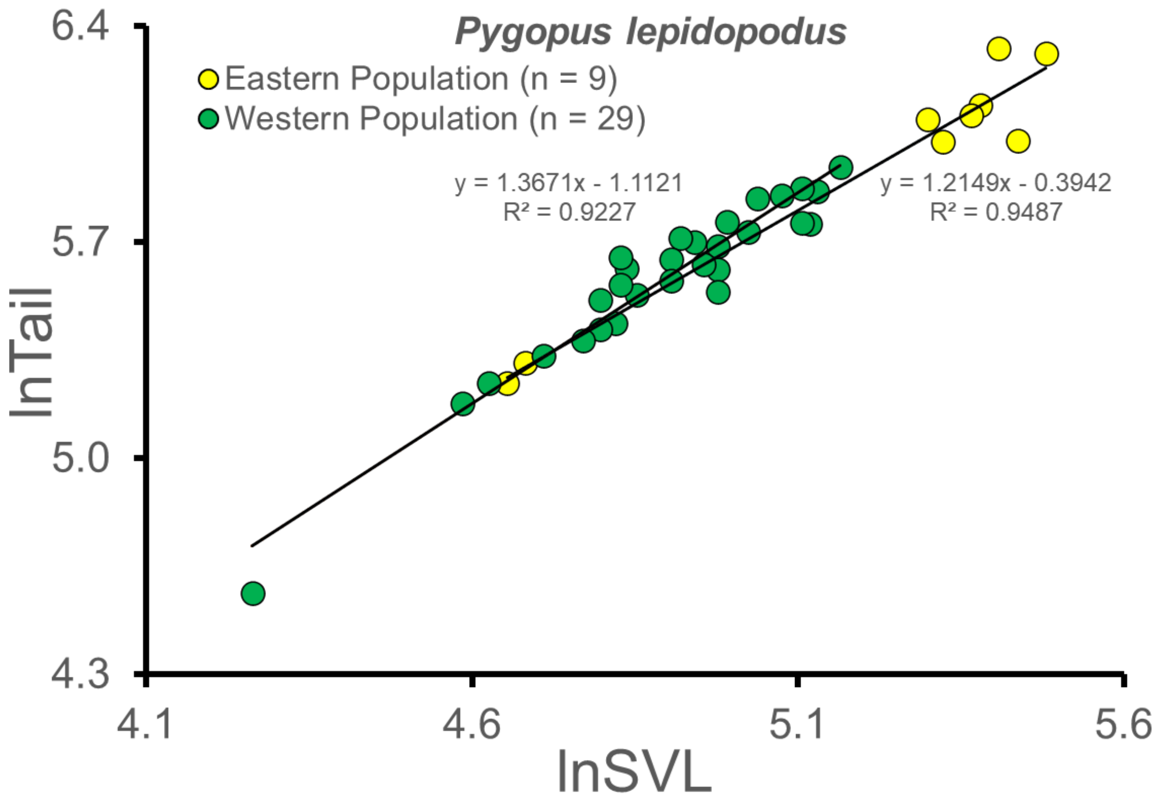


Supplementary File 1—Figure 1. Pairwise comparison of post-natal ontogenetic allometric trajectories for two populations of *Pygopus lepidopodus*. This plot shows the bivariate scaling relationships of ln-transformed tail length (ln*Tail*) on ln-transformed snout-vent length (ln*SVL*) for each population. Individuals from the Eastern Population (yellow circles) exhibited larger maximum body sizes (measured as maximum *SVL*) compared to individuals in the Western Population (green circles). The least squares regression equations and $R^2$ values are also shown. Note that the slopes of both trajectories are greater than 1.0 (i.e., isometry) and thus both trajectories appear to show positive ontogenetic allometry. Source data are available in Supplementary File 1—Table 1.

Supplementary File 1—Table 1. Source data for Supplementary File 1—Figure 1. All data for *Pygopus lepidopodus* were obtained from Jennings (2002). The "Eastern Population" refers to specimens from the states of New South Wales and Queensland, Australia, whereas the "Western Population" refers to specimens from the state of Western Australia. SVL = snout-vent length; Tail = tail length; AM = Australian Museum; QM = Queensland Museum; WAM = Western Australian Museum; and MCZ = Museum of Comparative Zoology, Harvard University; WBJ = W. Bryan Jennings field number.

| **Population** | **Specimen #** | **SVL (mm)** | **Tail (mm)** |
|---|---|---|---|
| *lepidopodus (Eastern population)* | WBJ 1837 (now MCZ R-187973) | 240 | 550 |
| *lepidopodus (Eastern Population)* | WBJ 2070 (now MCZ R-187943) | 230 | 415 |
| *lepidopodus (Eastern Population)* | WBJ 1864 (now MCZ R-187976) | 223 | 560 |
| *lepidopodus (Eastern Population)* | WBJ 2011 (now WAM R-128503) | 217 | 465 |
| *lepidopodus (Eastern Population)* | WBJ 1841 (now MCZ R-187975) | 214 | 450 |
| *lepidopodus (Eastern Population)* | WBJ 1840 (now MCZ R-187974) | 205 | 413 |
| *lepidopodus (Eastern Population)* | WBJ 2112 (now MCZ R-187945) | 200 | 445 |
| *lepidopodus (Eastern Population)* | WBJ 1241 (now WAM R-128477) | 108 | 202 |
| *lepidopodus (Eastern Population)* | WBJ 1565 (now MCZ R-187970) | 105 | 189 |
| *lepidopodus (Western Population)* | WBJ 1839 (now WAM R-128489) | 175 | 381 |
| *lepidopodus (Western Population)* | WBJ 1836 (now WAM R-128488) | 169 | 352 |
| *lepidopodus (Western Population)* | WBJ 1091 (now WAM R-128467) | 167 | 317 |
| *lepidopodus (Western Population)* | WBJ 1872 (now MCZ R-187977) | 165 | 356 |
| *lepidopodus (Western Population)* | WBJ 1236 (now MCZ R-187959) | 165 | 318 |
| *lepidopodus (Western Population)* | WBJ 1021 (now WAM R128466) | 160 | 347 |

| | | | |
|---|---|---|---|
| *lepidopodus (Western Population)* | WBJ 1147 (now WAM R-128471) | 154 | 344 |
| *lepidopodus (Western Population)* | WBJ 2471 (now MCZ R-187951) | 152 | 309 |
| *lepidopodus (Western Population)* | WBJ 1235 (now MCZ R-187963) | 147 | 319 |
| *lepidopodus (Western Population)* | WBJ 1418 (now MCZ R-187966) | 145 | 295 |
| *lepidopodus (Western Population)* | WBJ 1378 (now MCZ R-187964) | 145 | 273 |
| *lepidopodus (Western Population)* | WBJ 2312 (now MCZ R-187950) | 145 | 254 |
| *lepidopodus (Western Population)* | WBJ 2132 (now WAM R-128517) | 142 | 278 |
| *lepidopodus (Western Population)* | WBJ 2103 (now WAM R-128512) | 140 | 299 |
| *lepidopodus (Western Population)* | WBJ 2148 (now WAM R-128518) | 137 | 303 |
| *lepidopodus (Western Population)* | WBJ 1880 (now MCZ R-187938) | 135 | 283 |
| *lepidopodus (Western Population)* | WBJ 1889 (now WAM R-128500) | 135 | 264 |
| *lepidopodus (Western Population)* | WBJ 1099 (now MCZ R-187955) | 128 | 252 |
| *lepidopodus (Western Population)* | WBJ 1206 (now MCZ R-187957) | 126 | 274 |
| *lepidopodus (Western Population)* | WBJ 1810 (now MCZ R-187972) | 125 | 284 |
| *lepidopodus (Western Population)* | WBJ 1837 (now MCZ R-187973) | 125 | 260 |
| *lepidopodus (Western Population)* | WBJ 2070 (now MCZ R-187943) | 124 | 230 |
| *lepidopodus (Western Population)* | WBJ 1864 (now MCZ R-187976) | 121 | 248 |
| *lepidopodus (Western Population)* | WBJ 2011 (now WAM R-128503) | 121 | 225 |
| *lepidopodus (Western Population)* | WBJ 1841 (now MCZ R-187975) | 118 | 217 |
| *lepidopodus (Western Population)* | WBJ 1840 (now MCZ R-187974) | 111 | 207 |
| *lepidopodus (Western Population)* | WBJ 2112 (now MCZ R-187945) | 102 | 189 |
| *lepidopodus (Western Population)* | WBJ 1241 (now WAM R-128477) | 98 | 177 |
| *lepidopodus (Western Population)* | WBJ 1565 (now MCZ R-187970) | 71 | 96 |

### Using published data to identify interpopulational trait scaling patterns in pygopodid lizards

If a trait's ontogenetic or static allometry (OA) relationship is known for a group of closely related species at the level of species group or genus, then a simple test for geometric similarity (GS) can be applied to data consisting of only two quantities for each operational taxonomic unit or "OTU" (i.e., population, subspecies, or species). The first quantity is the maximum known body size or "maximum $X$ value" of each OTU, where $X$ is a measure of body size. The second quantity is the maximum relative trait size ratio or "maximum $Y/X$" value for each OTU, where $Y/X$ represents relative trait size for specific size-age class. Notice that if a positive OA trait exhibits a GS pattern, then the maximum $Y/X$ of the smaller OTU is expected to be equivalent to the maximum $Y/X$ value for the larger OTU (Figure 1A). If, on the other hand, a positive OA trait displays evolutionary allometry (EA) via ontogenetic scaling (Figure 1C) or EA via transposition patterns (Figure 1E), then the maximum $Y/X$ value for the larger OTU will be greater than the maximum $Y/X$ of the smaller OTU.

If a negative OA trait exhibits a GS pattern, then the maximum $Y/X$ of the smaller OTU is expected to be the same as the maximum $Y/X$ value for the larger OTU. Note, however, that in contrast to traits that exhibit positive OA, the maximum $Y/X$ for a trajectory showing negative ontogenetic allometry is found early in post-natal development. Therefore, juveniles will have larger $Y/X$ than adults along a given trajectory. However, this is not a complication provided that the $Y/X$ values for a pair of OTUs represent a comparable size-age class. If, on the other hand, a negative OA trait displays ontogenetic scaling (Figure 1D) or EA via transposition patterns

(Figure 1F), then the *Y*/*X* value for the smaller OTU will be greater than the corresponding *Y*/*X* of the larger OTU.

**Application of the test to published summary data for pygopodid lizards**

Pygopodid lizards exhibited a strong trend of positive ontogenetic allometry for tail length when the natural log of tail length or "ln*Tail*" was regressed against the natural log of snout-vent length or "ln*SVL*," a measure of body size (Jennings 2002). Using this information along with published estimates of maximum *SVL* and maximum relative tail length (maximum tail length as a percentage of *SVL*), it was possible to test several OTU pairs for GS using the above methodology. Because it was expected that very recently diverged conspecific populations (subspecies) or sister species would show tail divergence patterns consistent with geometric similarity, a search of the literature for appropriate pygopodid OTU pairs was conducted. The search yielded three pairs with the required data for this test. The first OTU pair (*Delma concinna concinna* vs. *D. c. major*) comprised two subspecies while the latter two pairs (*Delma haroldi* vs. *D. butleri* and *Pletholax gracilis* vs. *P. edelensis*) consisted of two very recently diverged sister species according to phylogenetic studies (Brennan et al 2016; Skipwith et al. 2019; Keally et al. 2020). For each pairwise comparison, the OTU with the smaller maximum body size showed a relatively longer tail (Supplementary File 1—Table 2). These results reject the hypothesis of positive EA while GS and negative EA cannot be ruled out. Accordingly, these results provided preliminary evidence suggesting that body size and tail shape had become dissociated in these lineages as would be expected for GS.

Supplementary File 1—Table 2. Published estimates of maximum snout-vent length (*SVL*) and maximum relative tail length (tail length expressed as percentage of *SVL*) for three pygopodid lizard OTU pairs. OTU = operational taxonomic unit (i.e., subspecies or species). Note that *Aclys concinna concinna* and *A. c. major* (vis-à-vis Storr 1987) are now referred to as *Delma concinna concinna and D. c. major* (Jennings et al. 2003), while *Pletholax gracilis edelensis* (vis-à-vis Storr 1978) is now *Pletholax edelensis* (Keally et al. 2020).

| pairwise comparison of comparable OTUs | OTU | maximum SVL (mm) | length of tail (% SVL) | source of data |
|---|---|---|---|---|
| *Delma concinna concinna* vs. *D. c. major* | *Delma concinna concinna* | 101 | 450% | Storr 1987 |
| *Delma concinna concinna* vs. *D. c. major* | *Delma concinna major* | 112 | 438% | Storr 1987 |
| *Delma haroldi* vs. *D. butleri* | *Delma haroldi* | 75 | 369% | Storr 1987 |
| *Delma haroldi* vs. *D. butleri* | *Delma butleri* | 91 | 345% | Storr 1987 |
| *Pletholax gracilis* vs. *P. edelensis* | *Pletholax gracilis* | 83 | 346% | Storr 1978 |
| *Pletholax gracilis* vs. *P. edelensis* | *Pletholax edelensis* | 90 | 340% | Storr 1978 |
