## Supplementary File 2 for "A scaling theory of trait evolution"

### Identifying interspecific trait scaling patterns between *Delma branchia* and *D. pax* of the Delma tincta species group using a combination of data from this study and a published study

Pepper et al. (2025) conducted a molecular and morphological study of the Delma tincta species group. We can use the testing framework described in Supplementary File 1 to determine if two members of this group, *D. branchia* and *D. pax*, which occur in sympatry and syntopy, exhibit patterns consistent with body size and trait shape dissociation (i.e., geometric similarity). To perform the test, we can use snout-vent-length (*SVL*), head width at the ears (*Head*W), and tail length (*TailL*) data for *D. branchia* in the Appendix 4 of Pepper et al. (2025) with comparable data for *D. pax* in this paper (see Supplementary Data and raw trait data). Note, the *HeadW* measure will be used as a proxy for average body width because body width data were not available for *D. branchia*.

These data suggest that *Delma pax* reaches a higher maximum *SVL* than *D. branchia* (Supplementary File 2—Table 1), consistent with Maryan et al. (2007), who called *D. branchia* by its former name, *D. tincta*. Results of tests for the three traits are shown in Supplementary File 2—Table 1. In the first two trait comparisons, *D. branchia* exhibits a longer relative tail length (using *SVL* and *HeadW* as the size references, respectively) compared to the larger *D. pax*. Given that *D. pax* exhibited positive ontogenetic allometry (OA) for tail length (see Supplementary Tables S1 and S2), which fits the trends in pygopodids, it is reasonable to assume that *D. branchia* also shows positive OA for these two traits. If this is correct, then these results concerning relative tail length are suggestive of geometric similarity (GS) or negative evolutionary allometry (EA). In contrast, the results of this study showed that relative body width shows negative OA for *D. pax*, which fits the trend in pygopodids (see Supplementary Table S3). If it is therefore assumed that *D. branchia* also exhibits negative OA for body width, then the *HeadW* on *SVL* scaling relationship reflects GS or positive EA. Unfortunately, it is not possible to distinguish GS from negative EA for the two tail length relationships, and GS from positive EA for the *HeadW* relationship, using these minimalistic data and analysis methods. Nonetheless, the results in Supplementary File 2—Table 1 suggest that the shapes of all three traits became dissociated from body size since the population divergence event that gave rise to the *D. pax* and *D. branchia* lineages, which is consistent with a geometric scaling scenario (Gould 1971).

Supplementary File 2—Table 1. Test for evolutionary dissociation between body size and trait shape between *Delma pax* and *D. branchia*. Maximum snout-vent length = maximum *SVL*, maximum tail length expressed as percentage of *SVL* = *TailL* (% *SVL*), maximum tail length expressed as a percentage of head width at the ear = *TailL* (% *HeadW*), and maximum head width at ear expressed as percentage of *SVL* = *HeadW* (% *SVL*).

| **Species** | **maximum *SVL* (mm)** | ***TailL* (% *SVL*)** | ***TailL* (% head width at ear)** | ***HeadW* (% *SVL*)** | **source of data** |
|---|---|---|---|---|---|
| *Delma pax* | 98 | 347% | 5567% | 8.8% | this study |
| *Delma branchia* | 93 | 368% | 5842% | 7.2% | Pepper et al. 2025 |
