## Supplementary Methods for "A scaling theory of trait evolution"



### Estimation of post-natal growth trajectories

Growth trajectories represented by the scaling relationship between a trait of interest and bodysize can be estimated using a linearized version of Huxley's (1932) allometric equation:

$$\ln Y = \alpha \ln X + \ln b \quad (1)$$

in which $\ln Y$ is the natural log-transformed trait variable ($Y$), $\ln X$ is the natural log-transformed body size variable ($X$), $\alpha$ is the slope, and $b$ is the $Y$-intercept. Following previous studies (e.g., Pounds et al., 1983; Garland, 1985), intraspecific samples of preserved lizards spanning the post-natal body-size spectrum were used in lieu of actual growth data. These "cross-sectional" data (Cock, 1966) were then used to generate an "ontogenetic" growth curve for each OTU. Although these data cannot be interpreted as individual growth trajectories, they may provide reasonable estimates of average ontogenetic growth within a population (Cock, 1966; Garland, 1985). The line-fitting method of reduced major axis (RMA) has often been employed for estimating allometric slopes owing to concerns that ordinary least squares (OLR) regression may underestimate the true slopes when measurement error exists in the data (e.g., Harvey and Pagel, 1991). However, others have recommended the use of OLR over RMA because the former is expected to perform well enough in most studies while the latter has drawbacks that make it unsuitable for allometry studies (Seim and Sæther 1983; Kelly and Price 2004; Hansen and Bartoszek 2012; Pélabon et al. 2014; Voje et al. 2014; Kilmer and Rodríguez 2017). Accordingly, ontogenetic allometric slopes were estimated via OLR using the *R* package *SMATR* (Warton and Weber, 2002; Warton et al. 2006; Taskinen and Warton 2011) implemented in the *R* statistical package (version 4.3.0., *R* Development Core Team, 2023).

One advantage to estimating linearized ontogenetic growth trajectories is that a test for isometry can be performed in which $\alpha$ is compared to a null isometric value (Sweet, 1980;

Pounds et al., 1983; Garland, 1985). When *Y* and *X* are the same dimensional units as they are in this study, an ontogenetic allometric regression with a slope of 1.0 indicates isometry, $\alpha > 1.0$ signifies positive ontogenetic allometry, and $\alpha < 1.0$ means negative ontogenetic allometry (Pounds et al. 1983; Garland 1985). Isometry tests were conducted using the *R* package *SMATR* and slopes were considered to be allometric if $p < 0.05$.

### Statistical analyses of pairwise comparisons of growth trajectories

Statistical tests using the *R* package *SMATR* were conducted to determine if each pair of ontogenetic trajectories overlapped each other (i.e., exhibited a common slope) or were transposed (i.e., showed different elevations). However, the test for common elevations is only meaningful if the test for common slopes indicates that the slopes are statistically the same. If the results indicated that a pair of slopes were equal, then a common elevations test based on the Wald's statistic (analogous to analysis of covariance) was performed. Null hypotheses of equal slopes and equal elevations were rejected if $p < 0.05$.

The eight scaling patterns were defined as follows: 1) *geometric similarity (all ages)* = statistical results indicated equal slopes and unequal elevations while visual analysis of ln-ln plots revealed that the two trajectories matched Figure 1A or 1B; 2) *evolutionary allometry overlap (equal slopes)* = statistical results indicated equal slopes and equal elevations while visual analysis revealed partially overlapping trajectories that matched Figure 1C or 1D—notice that this spatial pattern was simply defined as overlapping trajectories ("overlap") rather than ontogenetic scaling because the GSF model suggests that trajectories may partially overlap each other in coincidental fashion and thus cannot represent true ontogenetic scaling (see Phase 3 in Figure 2); 3) *evolutionary allometry transposition (all ages)* = statistical results indicated equal slopes and unequal elevations while visual analysis revealed that the two trajectories matched Figure 1E or 1F; 4) *geometric similarity (adults only*) = statistical analysis indicated unequal slopes while visual analysis revealed that the trajectories displayed a V-shaped pattern in which they overlapped for the juvenile classes while the adult classes diverged from each other in a transposition-like pattern that was consistent with Figure 1A or 1B; 5) *geometric similarity (juveniles only)* = statistical analysis indicated unequal slopes while visual analysis revealed that the trajectories displayed a V-shaped pattern in which they overlapped for the adult classes while the juvenile classes diverged from each other in a transposition-like pattern that was consistent with Figure 1A or 1B; 6) *evolutionary allometry overlap (unequal slopes)* = statistical results indicated unequal slopes while visual analysis revealed partially overlapping trajectories that formed a flattened X-shaped pattern; 7) *evolutionary allometry transposition (adults only)* = statistical analysis indicated unequal slopes while visual analysis revealed that the trajectories displayed a V-shaped pattern in which they overlapped for the juvenile classes while the adult classes diverged from each other in a transposition-like pattern that was consistent with Figure 1E or 1F; and 8) *evolutionary allometry transposition (juveniles only)* = statistical analysis indicated unequal slopes while visual analysis revealed that the trajectories displayed a V-shaped pattern in which they overlapped for the adult classes while the juvenile classes diverged from each other in a transposition-like pattern that was consistent with Figure 1E or 1F.

Note that scaling patterns 1-3 were straightforward to identify using a combination of statistical results and unambiguous graphical scaling patterns of regression lines on ln-ln plots while scaling patterns 4-8 were identified using statistical results and subjective determinations of graphical scaling patterns. Although the aforementioned statistical procedures can objectively discriminate between overlapping trajectories and transpositions that reflect geometric similarity or evolutionary allometry, intermediate patterns between these extremes may exist. If the GSF model is correct, then we could expect to see many such intermediates, which would make it more challenging to unambiguously identify scaling patterns. This may mean that it is not possible to eliminate all subjectivity from these analyses.
