## Supplementary Figures S1-S7 for "A scaling theory of trait evolution"

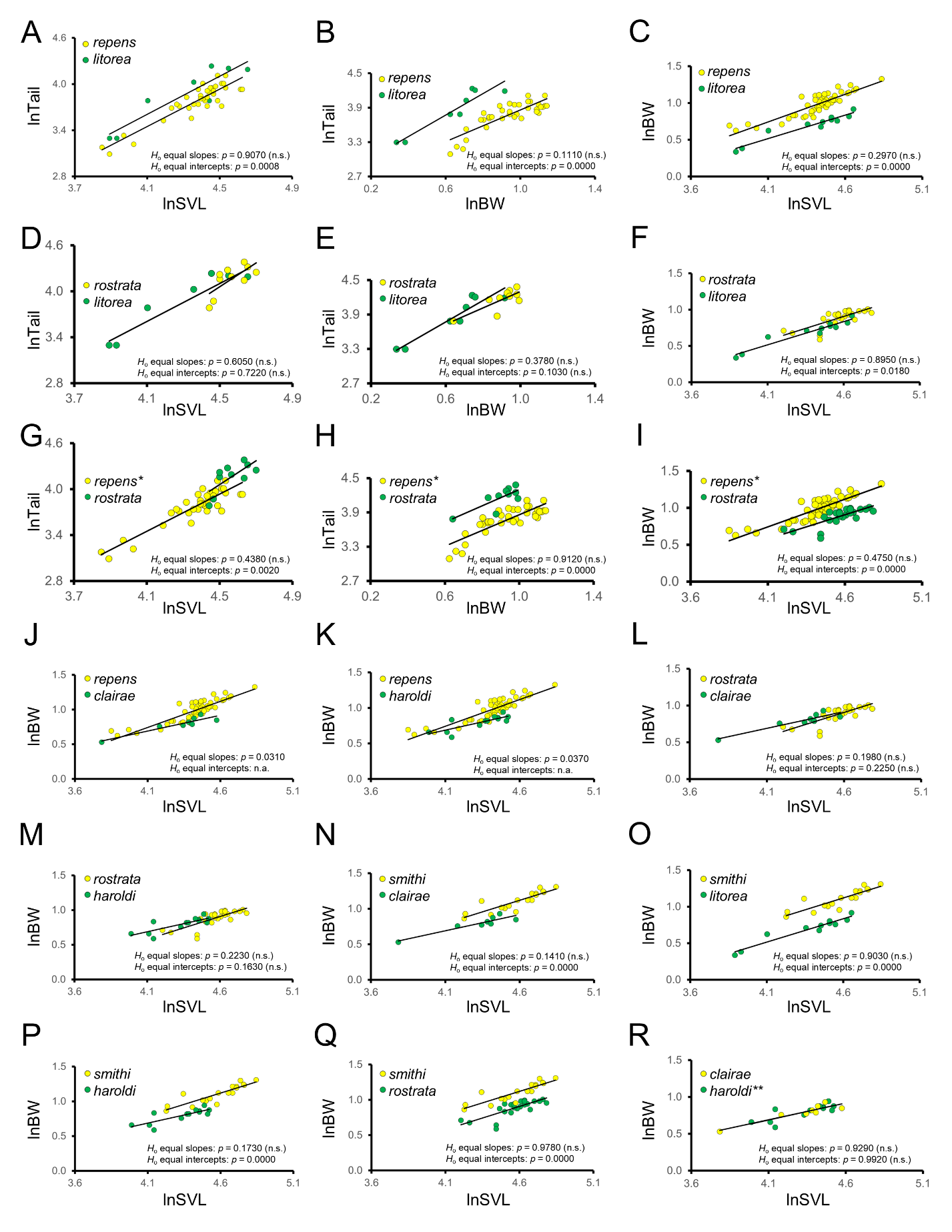

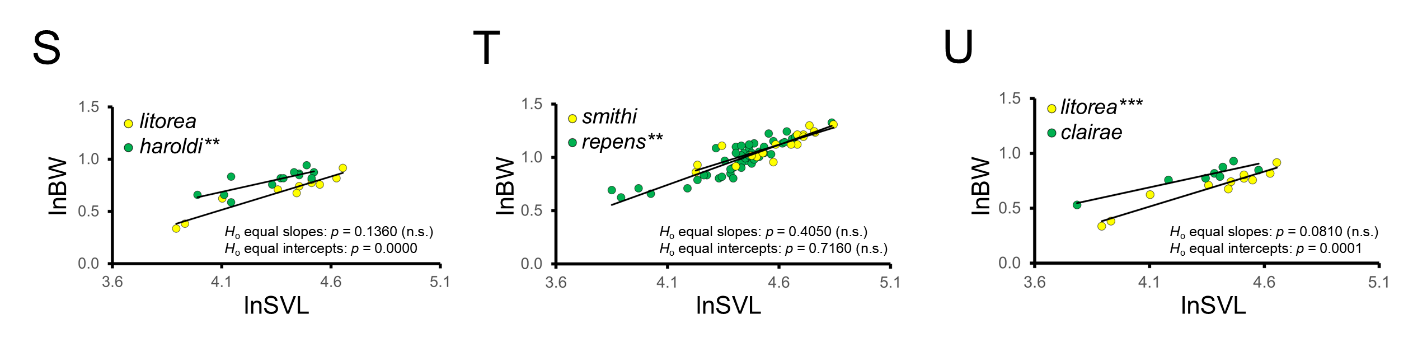
Figure S1. Pairwise comparisons of post-natal ontogenetic trajectories for the *Aprasia repens* species group. Plots A-I show bivariate scaling relationships ln*Tail* on ln*SVL*, ln*Tail* on ln$\bar{BW}$, and ln$\bar{BW}\mathrm{on}$ln*SVL* for the *A. repens*/*A. litorea*, *A. rostrata*/*A. litorea*, and *A. repens*/*A. rostrata* pairs. Plots J-U only show the ln$\bar{BW}\mathrm{on}$ln*SVL* relationship for other OTU pairs because one or both OTUs had insufficient tail length data (i.e., due to regenerated or incomplete tails on many specimens). In plots A-F and J-U, yellow OTUs exhibited longer maximum *SVL* compared to corresponding green OTUs. In A-Q, yellow OTUs had a larger maximum $\bar{BW}$ than corresponding green OTUs. In G-I, both OTUs have the same maximum *SVL* but the yellow OTU (*) had a larger maximum $\bar{BW}$ than the green OTU. In R-T, green OTUs (**) had larger maximum $\bar{BW}$ compared to corresponding yellow OTUs. In plot U, the yellow OTU (***) had a longer maximum *SVL* than the green OTU but both OTUs had the same maximum $\bar{BW}$. Each dot represents an individual. Results of the equal slopes and equal *Y*-intercepts tests are shown. *p-*values > 0.05 were statistically non-significant (n.s.). Equal intercepts tests were not applicable (n.a.) whenever equal slopes hypotheses were rejected. See Supplementary Data for maximum *SVL* and $\bar{BW}$ values and Supplementary Table S4 for a summary of scaling patterns obtained from this figure.

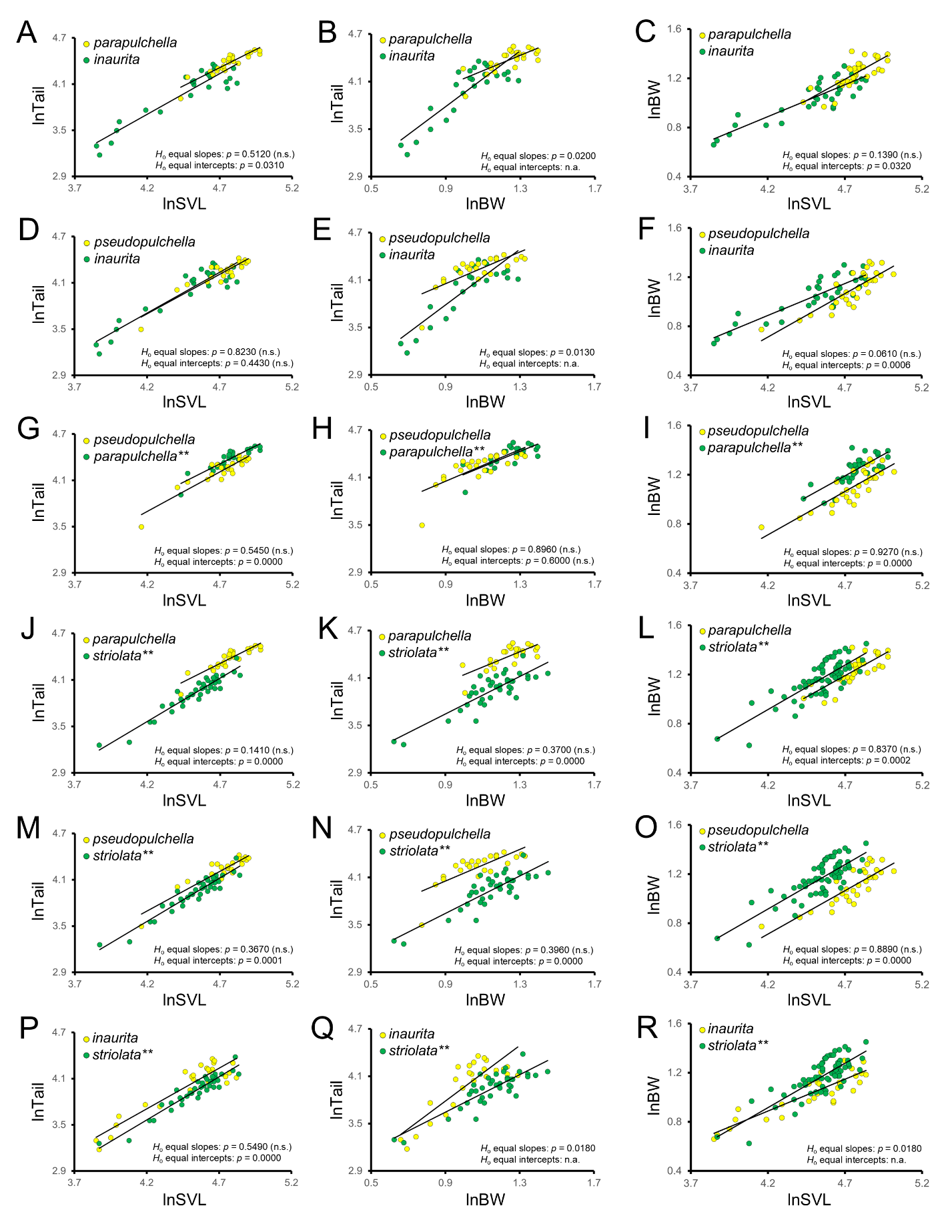

Figure S2. Pairwise comparisons of post-natal ontogenetic trajectories for the southeastern Australia *Aprasia* species group. Plots A-R show bivariate scaling relationships ln*Tail* on ln*SVL*, ln*Tail* on ln$\bar{BW}$, and ln$\bar{BW}\mathrm{on}$ln*SVL* comparisons for each OTU pair. In all plots, yellow OTUs exhibited longer maximum *SVL* compared to corresponding green OTUs. In A-F, yellow OTUs had larger maximum $\bar{BW}$ compared to corresponding green OTUs, whereas in G-R green OTUs (**) had larger maximum $\bar{BW}$ than corresponding yellow OTUs. Each dot represents an individual. Results of the equal slopes and equal *Y*-intercepts tests are shown. *p-*values > 0.05 were statistically non-significant (n.s.). Equal intercepts tests were not applicable (n.a.) whenever equal slopes hypotheses were rejected. See Supplementary Data for maximum *SVL* and $\bar{BW}$ values and Supplementary Table S5 for a summary of scaling patterns obtained from this figure.

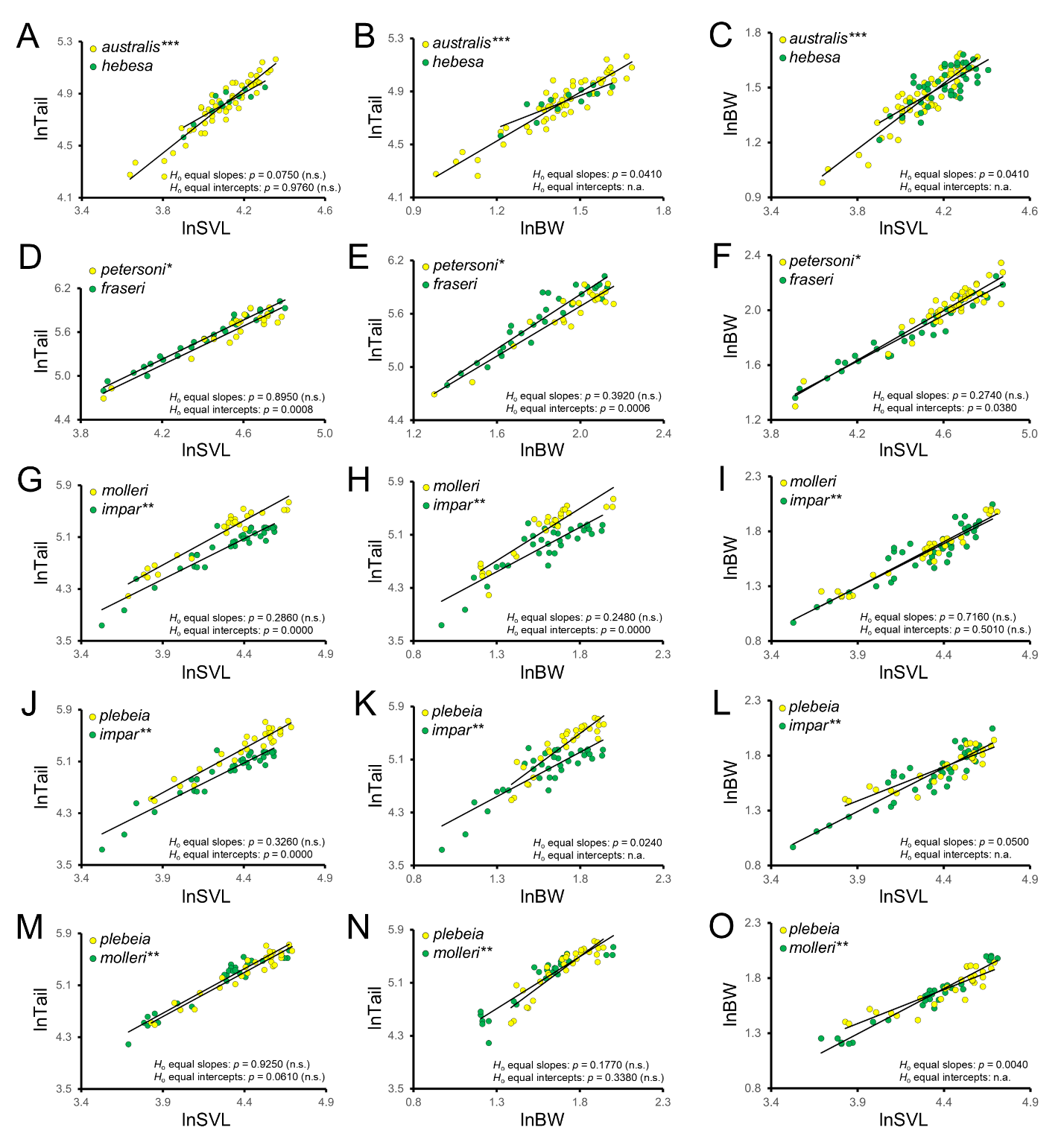
Figure S3. Pairwise comparisons of post-natal ontogenetic trajectories for OTUs in the *Delma australis*, *D. fraseri*, and *D. impar* species groups. Plots A-C show bivariate scaling relationships ln*Tail* on ln*SVL*, ln*Tail* on ln$\bar{BW}$, and ln$\bar{BW}\mathrm{on}$ln*SVL* for the *D. australis*/*D. hebesa* pair, plots D-F exhibit scaling relationships for the *D. petersoni*/*D. fraseri* pair, and plots G-O display scaling relationships for the *D. impar* group pairs. In A-C, the yellow OTU (***) had a longer maximum *SVL* compared to the green OTU, but both OTUs had the same maximum $\bar{BW}$. In D-F, both OTUs had the same maximum *SVL* but the yellow OTU (*) had a larger maximum $\bar{BW}$ compared to the green OTU. In G-O, yellow OTUs had longer maximum *SVL* compared to corresponding green OTUs while green OTUs (**) had larger maximum $\bar{BW}$ than corresponding yellow OTUs. Each dot represents an individual. Results of the equal slopes and equal *Y*-intercepts tests are shown. *p-*values > 0.05 were statistically non-significant (n.s.). Equal intercepts tests were not applicable (n.a.) whenever equal slopes hypotheses were rejected. See Supplementary Data for maximum *SVL* and $\bar{BW}$ values and Supplementary Table S6 for a summary of scaling patterns obtained from this figure.

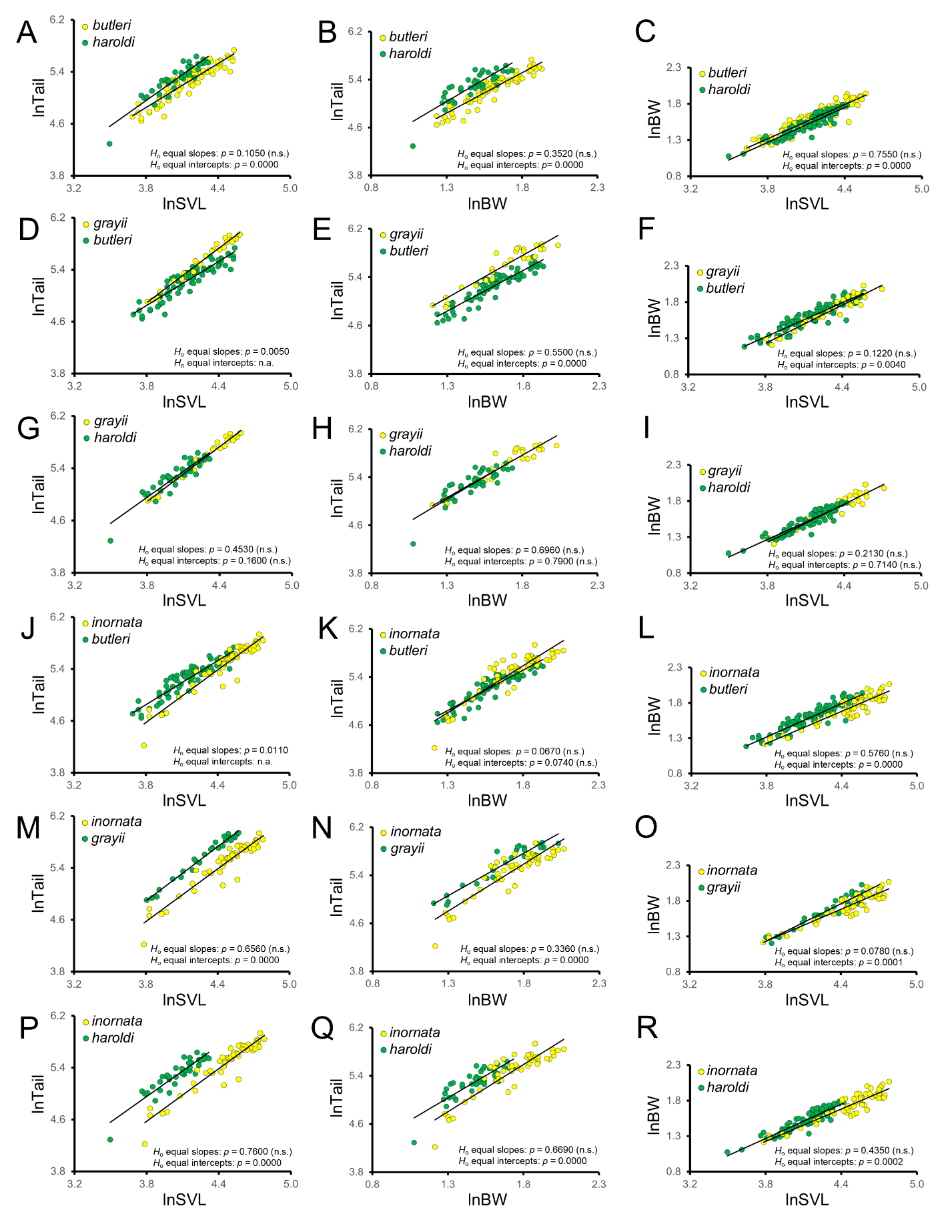

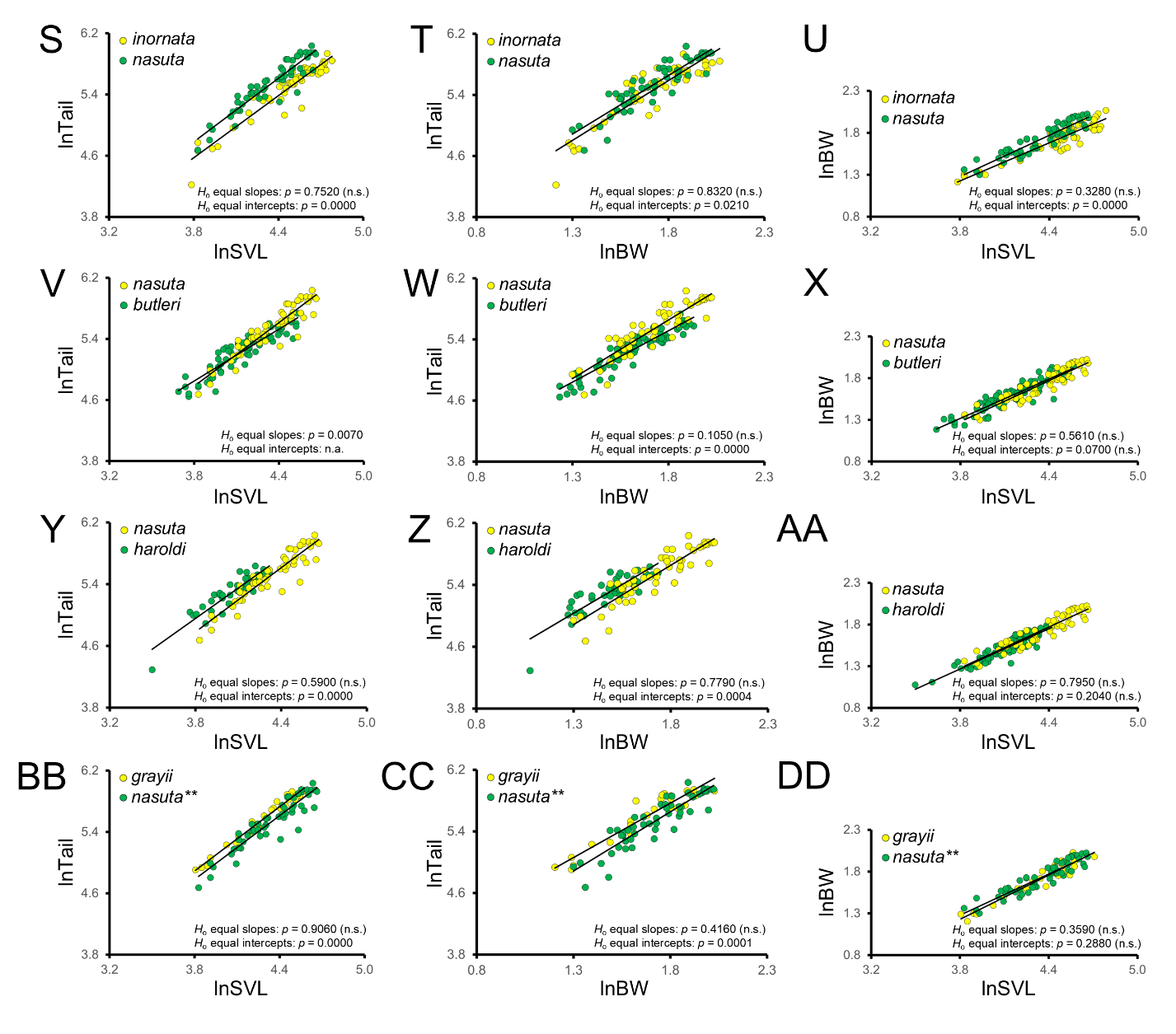

Figure S4. Pairwise comparisons of post-natal ontogenetic trajectories for the *Delma nasuta* species group. Plots A-DD show bivariate scaling relationships ln*Tail* on ln*SVL*, ln*Tail* on ln$\bar{BW}$, and ln$\bar{BW}\mathrm{on}$ln*SVL* for each OTU pair. In all plots, yellow OTUs exhibited longer maximum *SVL* compared to corresponding green OTUs. In A-AA, yellow OTUs had larger maximum $\bar{BW}$ compared to corresponding green OTUs. In BB-DD, the green OTU (**) had a larger maximum $\bar{BW}$ than the yellow OTU. Each dot represents an individual. Results of the equal slopes and equal *Y*-intercepts tests are shown. *p-*values > 0.05 were statistically non-significant (n.s.). Equal intercepts tests were not applicable (n.a.) whenever equal slopes hypotheses were rejected. See Supplementary Data for maximum *SVL* and $\bar{BW}$ values and Supplementary Table S7 for a summary of scaling patterns obtained from this figure.

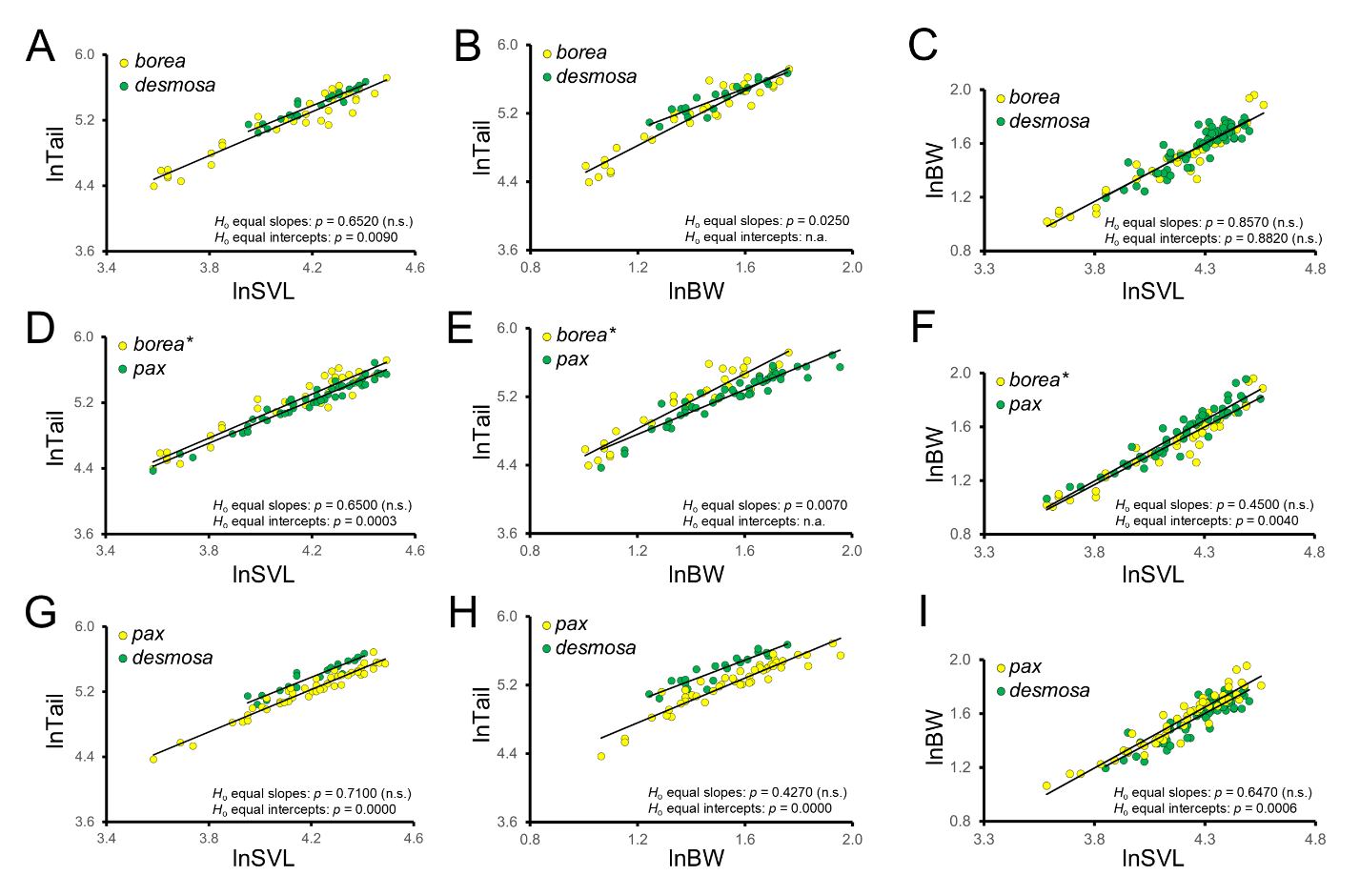
Figure S5. Pairwise comparisons of post-natal ontogenetic trajectories for the *Delma tincta* species group. Plots A-I show bivariate scaling relationships ln*Tail* on ln*SVL*, ln*Tail* on ln$\bar{BW}$, and ln$\bar{BW}\mathrm{on}$ln*SVL* for each OTU pair. In plots A-C and G-I, yellow OTUs exhibited longer maximum *SVL* and larger maximum $\bar{BW}$ than corresponding green OTUs, whereas in D-F both OTUs had the same maximum *SVL* but the yellow OTU (*) had a larger maximum $\bar{BW}$ compared to the green OTU. Each dot represents an individual. Results of the equal slopes and equal *Y*-intercepts tests are shown. *p-*values > 0.05 were statistically non-significant (n.s.). Equal intercepts tests were not applicable (n.a.) whenever equal slopes hypotheses were rejected. See Supplementary Data for maximum *SVL* and $\bar{BW}$ values and Supplementary Table S8 for a summary of scaling patterns obtained from this figure.

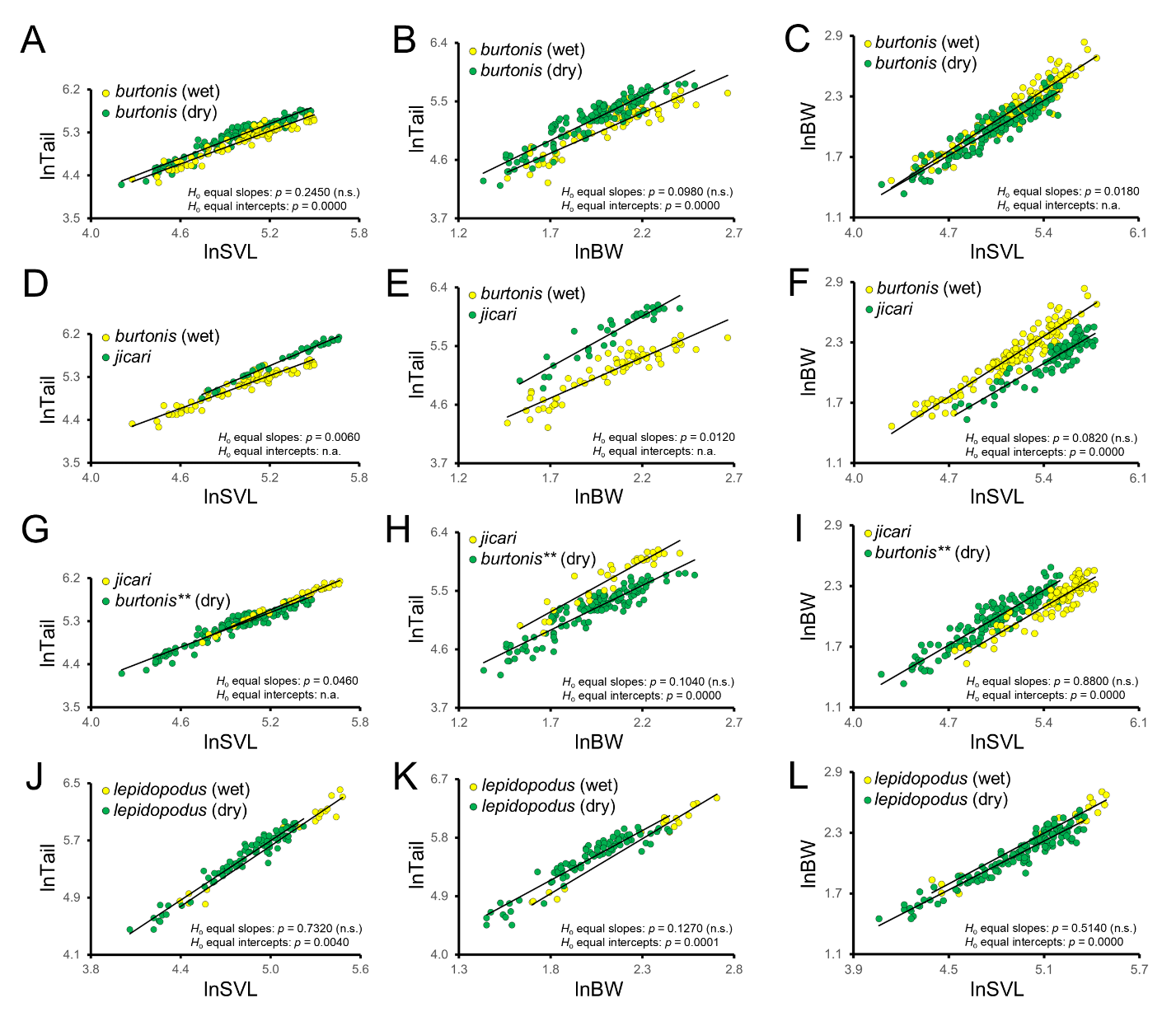
Figure S6. Pairwise comparisons of post-natal ontogenetic trajectories for the *Lialis* species group and two populations of *Pygopus lepidopodus*. Plots A-I show bivariate scaling relationships ln*Tail* on ln*SVL*, ln*Tail* on ln$\bar{BW}$, and ln$\bar{BW}\mathrm{on}$ln*SVL* for OTUs of the *Lialis* group, whereas plots J-L exhibit scaling relationships between two populations of *P. lepidopodus*. In all plots, yellow OTUs exhibited longer maximum *SVL* compared to corresponding green OTUs. In A-F and J-L, yellow OTUs had larger maximum $\bar{BW}$ compared to corresponding green OTUs. In G-I, the green OTU (**) had a larger maximum $\bar{BW}$ than the yellow OTU. Each dot represents an individua and “wet” and “dry” refer to *P. lepidopodus* populations that live in high and low rainfall biomes, respectively. Results of the equal slopes and equal *Y*-intercepts tests are shown. *p-*values > 0.05 were statistically non-significant (n.s.). Equal intercepts tests were not applicable (n.a.) whenever equal slopes hypotheses were rejected. See Supplementary Data for maximum *SVL* and $\bar{BW}$ values and Supplementary Table S9 for a summary of scaling patterns obtained from this figure.

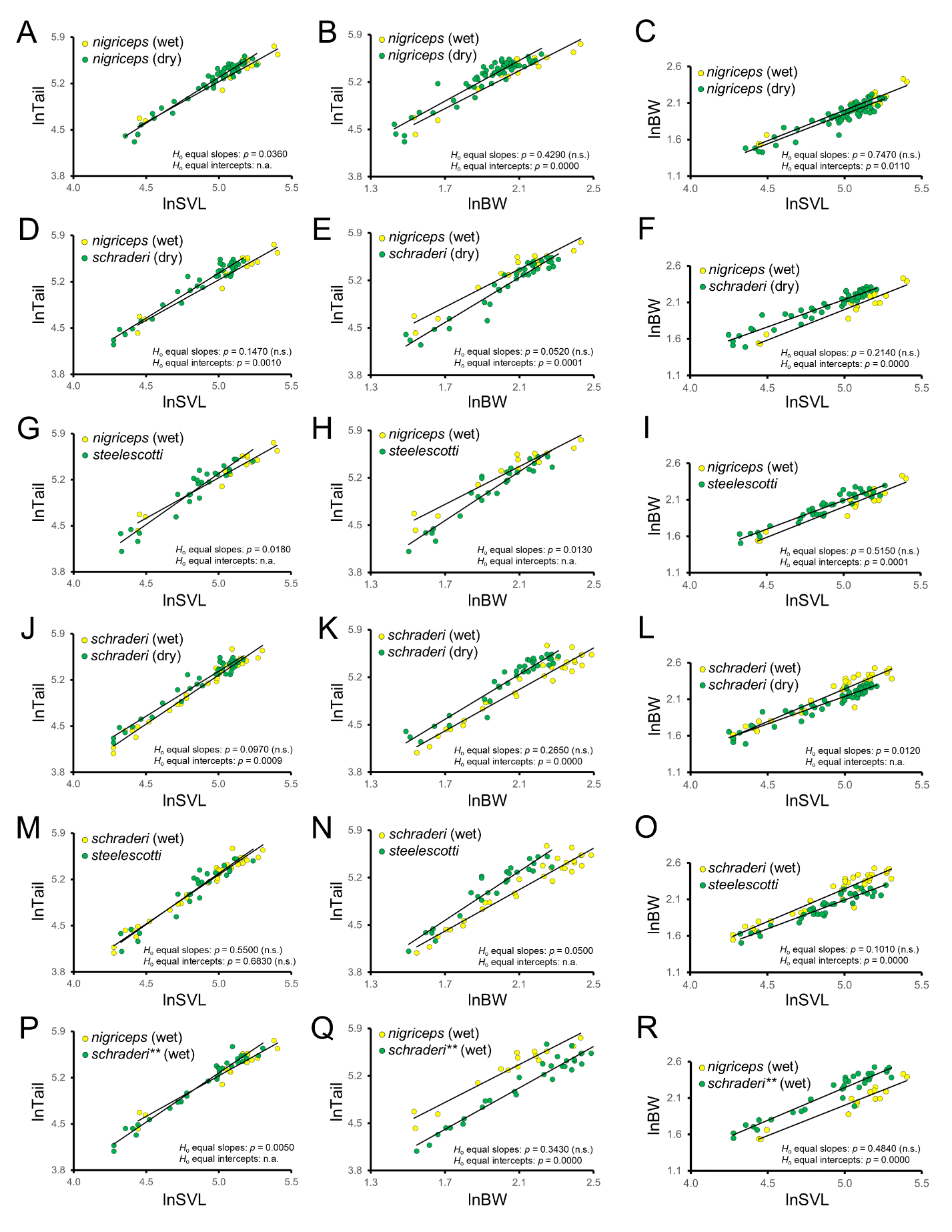

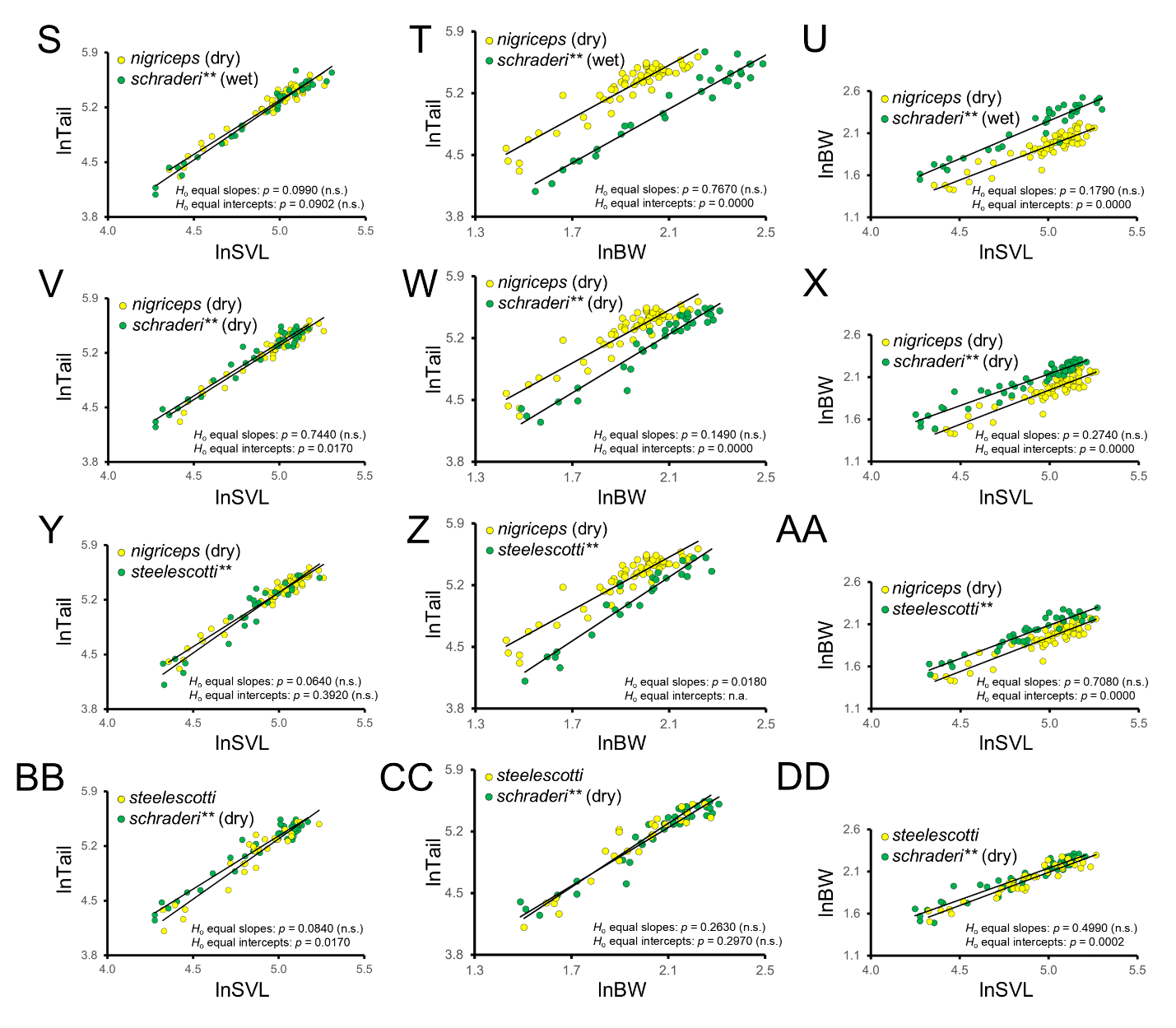
Figure S7. Pairwise comparisons of post-natal ontogenetic trajectories for the *Pygopus nigriceps* species groups. Plots A-DD show bivariate scaling relationships ln*Tail* on ln*SVL*, ln*Tail* on ln$\bar{BW}$, and ln$\bar{BW}\mathrm{on}$ln*SVL* for OTUs of the *P. nigriceps* group. In all plots, yellow OTUs exhibited longer maximum *SVL* compared to corresponding green OTUs. In A-O, yellow OTUs had larger maximum $\bar{BW}$ compared to corresponding green OTUs. In P-DD, green OTUs (**) had larger maximum $\bar{BW}$ than corresponding yellow OTUs. Each dot represents an individual and “wet” and “dry” refer to populations that live in high and low rainfall biomes, respectively. Results of the equal slopes and equal *Y*-intercepts tests are shown. *p-*values > 0.05 were statistically non-significant (n.s.). Equal intercepts tests were not applicable (n.a.) whenever equal slopes hypotheses were rejected. See Supplementary Data for maximum *SVL* and $\bar{BW}$ values and Supplementary Table S10 for a summary of scaling patterns obtained from this figure.
