## Supplementary Tables S1-S11 for "A scaling theory of trait evolution"



Table S1. Ontogenetic isometry tests for the ln*Tail* on ln*SVL* relationship in pygopodid lizards. Slopes are negatively or positively allometric when they are significantly smaller or larger, respectively, than the null value of 1.0 (isometry). n = subsample size for each OTU; slope = ordinary least squares regression slope; 95% C.I. = 95% confidence interval; and $R^2$ = coefficient of determination. *p*-values shaded in gray are statistically significant at the 0.05 level.

| Species/Population Group | OTU | n | Slope | 95% C.I. | $R^2$ | *p*-value |
|---|---|---|---|---|---|---|
| Aprasia repens group | *Aprasia litorea* | 8 | 1.24 | (0.70-1.77) | 0.84 | 0.3184 |
| Aprasia repens group | *Aprasia repens* | 37 | 1.21 | (1.02-1.41) | 0.82 | 0.0320 |
| Aprasia repens group | *Aprasia rostrata* | 10 | 1.56 | (0.34-2.78) | 0.52 | 0.3235 |
| Southeastern Australia Aprasia group | *Aprasia inaurita* | 26 | 1.06 | (0.88-1.23) | 0.86 | 0.5212 |
| Southeastern Australia Aprasia group | *Aprasia parapulchella* | 25 | 0.94 | (0.75-1.14) | 0.81 | 0.5480 |
| Southeastern Australia Aprasia group | *Aprasia pseudopulchella* | 25 | 1.02 | (0.84-1.21) | 0.85 | 0.7951 |
| Southeastern Australia Aprasia group | *Aprasia striolata* | 39 | 1.12 | (1.00-1.24) | 0.91 | 0.0516 |
| Delma australis group | *Delma australis* | 53 | 1.24 | (1.11-1.36) | 0.88 | 0.0005 |
| Delma australis group | *Delma hebesa* | 13 | 0.88 | (0.57-1.19) | 0.78 | 0.4181 |
| Delma fraseri group | *Delma fraseri* | 30 | 1.35 | (1.26-1.44) | 0.97 | 0.0000 |
| Delma fraseri group | *Delma petersoni* | 23 | 1.33 | (1.13-1.54) | 0.90 | 0.0026 |
| Delma impar group | *Delma impar* | 36 | 1.25 | (1.09-1.41) | 0.88 | 0.0026 |
| Delma impar group | *Delma molleri* | 26 | 1.37 | (1.21-1.54) | 0.93 | 0.0001 |
| Delma impar group | *Delma plebeia* | 29 | 1.36 | (1.21-1.52) | 0.93 | 0.0000 |
| Delma nasuta group | *Delma butleri* | 73 | 1.13 | (1.02-1.25) | 0.85 | 0.0259 |
| Delma nasuta group | *Delma grayii* | 27 | 1.40 | (1.31-1.48) | 0.98 | 0.0000 |
| Delma nasuta group | *Delma haroldi* | 37 | 1.32 | (1.12-1.52) | 0.84 | 0.0029 |
| Delma nasuta group | *Delma inornata* | 50 | 1.35 | (1.22-1.49) | 0.90 | 0.0000 |
| Delma nasuta group | *Delma nasuta* | 52 | 1.38 | (1.23-1.54) | 0.87 | 0.0000 |
| Delma tincta group | *Delma borea* | 34 | 1.34 | (1.19-1.49) | 0.92 | 0.0000 |
| Delma tincta group | *Delma desmosa* | 21 | 1.26 | (1.07-1.46) | 0.91 | 0.0110 |
| Delma tincta group | *Delma pax* | 49 | 1.30 | (1.22-1.39) | 0.95 | 0.0000 |
| Lialis group | *Lialis burtonis* (wet biome) | 71 | 1.13 | (1.04-1.21) | 0.92 | 0.0031 |
| Lialis group | *Lialis burtonis* (dry biome) | 130 | 1.21 | (1.15-1.27) | 0.93 | 0.0000 |
| Lialis group | *Lialis jicari* | 34 | 1.33 | (1.27-1.39) | 0.99 | 0.0000 |
| Pygopus lepidopodus group | *Pygopus lepidopodus* (wet biome) | 18 | 1.41 | (1.28-1.54) | 0.97 | 0.0000 |
| Pygopus lepidopodus group | *Pygopus lepidopodus* (dry biome) | 84 | 1.39 | (1.32-1.46) | 0.95 | 0.0000 |
| Pygopus nigriceps group | *Pygopus nigriceps* (wet biome) | 15 | 1.23 | (1.06-1.39) | 0.95 | 0.0113 |
| Pygopus nigriceps group | *Pygopus nigriceps* (dry biome) | 62 | 1.38 | (1.30-1.46) | 0.95 | 0.0000 |
| Pygopus nigriceps group | *Pygopus schraderi* (wet biome) | 31 | 1.48 | (1.39-1.58) | 0.97 | 0.0000 |
| Pygopus nigriceps group | *Pygopus schraderi* (dry biome) | 36 | 1.36 | (1.25-1.47) | 0.95 | 0.0000 |
| Pygopus nigriceps group | *Pygopus steelescotti* | 27 | 1.55 | (1.35-1.74) | 0.91 | 0.0000 |

Table S2. Ontogenetic isometry tests for the ln$Tail$ on ln$\overline{BW}$ relationship in pygopodid lizards. Slopes are negatively or positively allometric when they are significantly smaller or larger, respectively, than the null value of 1.0 (isometry). n = subsample size for each OTU; slope = ordinary least squares regression slope; 95% C.I. = 95% confidence interval; and $R^2$ = coefficient of determination. $p$-values shaded in gray are statistically significant at the 0.05 level.

| Species/Population Group | OTU | n | Slope | 95% C.I. | $R^2$ | $p$-value |
|---|---|---|---|---|---|---|
| Aprasia repens group | *Aprasia litorea* | 8 | 1.87 | (1.22-2.53) | 0.89 | 0.0172 |
| Aprasia repens group | *Aprasia repens* | 37 | 1.38 | (1.07-1.69) | 0.70 | 0.0173 |
| Aprasia repens group | *Aprasia rostrata* | 10 | 1.43 | (0.51-2.35) | 0.62 | 0.3149 |
| Southeastern Australia Aprasia group | *Aprasia inaurita* | 26 | 1.76 | (1.30-2.21) | 0.73 | 0.0021 |
| Southeastern Australia Aprasia group | *Aprasia parapulchella* | 25 | 0.97 | (0.61-1.33) | 0.57 | 0.8636 |
| Southeastern Australia Aprasia group | *Aprasia pseudopulchella* | 25 | 1.00 | (0.64-1.37) | 0.58 | 0.9843 |
| Southeastern Australia Aprasia group | *Aprasia striolata* | 39 | 1.19 | (0.94-1.44) | 0.72 | 0.1342 |
| Delma australis group | *Delma australis* | 53 | 1.23 | (1.09-1.36) | 0.87 | 0.0015 |
| Delma australis group | *Delma hebesa* | 13 | 0.81 | (0.50-1.13) | 0.75 | 0.2155 |
| Delma fraseri group | *Delma fraseri* | 30 | 1.53 | (1.33-1.74) | 0.89 | 0.0000 |
| Delma fraseri group | *Delma petersoni* | 23 | 1.40 | (1.18-1.63) | 0.89 | 0.0012 |
| Delma impar group | *Delma impar* | 36 | 1.34 | (1.06-1.62) | 0.73 | 0.0200 |
| Delma impar group | *Delma molleri* | 26 | 1.57 | (1.29-1.85) | 0.85 | 0.0003 |
| Delma impar group | *Delma plebeia* | 29 | 1.86 | (1.53-2.20) | 0.83 | 0.0000 |
| Delma nasuta group | *Delma butleri* | 73 | 1.35 | (1.22-3.30) | 0.86 | 0.0000 |
| Delma nasuta group | *Delma grayii* | 27 | 1.41 | (1.22-1.61) | 0.90 | 0.0002 |
| Delma nasuta group | *Delma haroldi* | 37 | 1.48 | (1.17-1.79) | 0.73 | 0.0031 |
| Delma nasuta group | *Delma inornata* | 50 | 1.57 | (1.35-1.78) | 0.81 | 0.0000 |
| Delma nasuta group | *Delma nasuta* | 52 | 1.54 | (1.33-1.74) | 0.82 | 0.0000 |
| Delma tincta group | *Delma borea* | 34 | 1.61 | (1.42-1.79) | 0.91 | 0.0000 |
| Delma tincta group | *Delma desmosa* | 21 | 1.19 | (0.96-1.41) | 0.86 | 0.0996 |
| Delma tincta group | *Delma pax* | 49 | 1.31 | (1.17-1.44) | 0.89 | 0.0000 |
| Lialis group | *Lialis burtonis* (wet biome) | 72 | 1.21 | (1.08-1.33) | 0.84 | 0.0018 |
| Lialis group | *Lialis burtonis* (dry biome) | 130 | 1.37 | (1.28-1.47) | 0.86 | 0.0000 |
| Lialis group | *Lialis jicari* | 34 | 1.56 | (1.35-1.76) | 0.88 | 0.0000 |
| Pygopus lepidopodus group | *Pygopus lepidopodus* (wet biome) | 18 | 1.69 | (1.55-1.83) | 0.98 | 0.0000 |
| Pygopus lepidopodus group | *Pygopus lepidopodus* (dry biome) | 84 | 1.53 | (1.42-1.65) | 0.90 | 0.0000 |
| Pygopus nigriceps group | *Pygopus nigriceps* (wet biome) | 15 | 1.40 | (1.18-1.62) | 0.94 | 0.0017 |
| Pygopus nigriceps group | *Pygopus nigriceps* (dry biome) | 62 | 1.50 | (1.35-1.65) | 0.88 | 0.0000 |
| Pygopus nigriceps group | *Pygopus schraderi* (wet biome) | 31 | 1.53 | (1.37-1.69) | 0.93 | 0.0000 |
| Pygopus nigriceps group | *Pygopus schraderi* (dry biome) | 36 | 1.66 | (1.50-1.82) | 0.93 | 0.0000 |
| Pygopus nigriceps group | *Pygopus steelescotti* | 27 | 1.81 | (1.57-2.50) | 0.91 | 0.0000 |

Table S3. Ontogenetic isometry tests for the ln$\overline{BW}$ on ln$SVL$ relationship in pygopodid lizards. Slopes are negatively or positively allometric when they are significantly smaller or larger, respectively, than the null value of 1.0 (isometry). n = subsample size for each OTU; slope = ordinary least squares regression slope; 95% C.I. = 95% confidence interval; and $R^2$ = coefficient of determination. $p$-values shaded in gray are statistically significant at the 0.05 level.

| Species/Population Group | OTU | n | Slope | 95% C.I. | $R^2$ | $p$-value |
|---|---|---|---|---|---|---|
| Aprasia repens group | *Aprasia clairae* | 8 | 0.45 | (0.27-0.64) | 0.86 | 0.0003 |
| Aprasia repens group | *Aprasia haroldi* | 14 | 0.47 | (0.26-0.67) | 0.67 | 0.0001 |
| Aprasia repens group | *Aprasia litorea* | 11 | 0.64 | (0.50-0.78) | 0.92 | 0.0002 |
| Aprasia repens group | *Aprasia repens* | 48 | 0.75 | (0.63-0.88) | 0.76 | 0.0002 |
| Aprasia repens group | *Aprasia rostrata* | 25 | 0.66 | (0.42-0.89) | 0.59 | 0.0061 |
| Aprasia repens group | *Aprasia smithi* | 18 | 0.65 | (0.46-0.84) | 0.76 | 0.0014 |
| Southeastern Australia Aprasia group | *Aprasia inaurita* | 34 | 0.52 | (0.42-0.62) | 0.77 | 0.0000 |
| Southeastern Australia Aprasia group | *Aprasia parapulchella* | 33 | 0.70 | (0.49-0.90) | 0.61 | 0.0050 |
| Southeastern Australia Aprasia group | *Aprasia pseudopulchella* | 31 | 0.71 | (0.53-0.88) | 0.71 | 0.0017 |
| Southeastern Australia Aprasia group | *Aprasia striolata* | 69 | 0.72 | (0.59-0.86) | 0.64 | 0.0000 |
| Delma australis group | *Delma australis* | 56 | 0.90 | (0.79-1.01) | 0.83 | 0.0662 |
| Delma australis group | *Delma hebesa* | 43 | 0.68 | (0.50-0.87) | 0.57 | 0.0012 |
| Delma fraseri group | *Delma fraseri* | 32 | 0.83 | (0.76-0.91) | 0.94 | 0.0001 |
| Delma fraseri group | *Delma petersoni* | 45 | 0.90 | (0.80-1.00) | 0.89 | 0.0530 |
| Delma impar group | *Delma impar* | 38 | 0.79 | (0.67-0.92) | 0.83 | 0.0016 |
| Delma impar group | *Delma molleri* | 29 | 0.82 | (0.75-0.90) | 0.95 | 0.0001 |
| Delma impar group | *Delma plebeia* | 30 | 0.62 | (0.49-0.74) | 0.79 | 0.0000 |
| Delma nasuta group | *Delma butleri* | 93 | 0.79 | (0.72-0.85) | 0.86 | 0.0000 |
| Delma nasuta group | *Delma grayii* | 29 | 0.88 | (0.78-0.98) | 0.92 | 0.0253 |
| Delma nasuta group | *Delma haroldi* | 74 | 0.80 | (0.72-0.89) | 0.85 | 0.0000 |
| Delma nasuta group | *Delma inornata* | 54 | 0.76 | (0.67-0.85) | 0.85 | 0.0000 |
| Delma nasuta group | *Delma nasuta* | 53 | 0.82 | (0.73-0.91) | 0.87 | 0.0002 |
| Delma tincta group | *Delma borea* | 38 | 0.86 | (0.77-0.95) | 0.91 | 0.0027 |
| Delma tincta group | *Delma desmosa* | 64 | 0.87 | (0.76-0.99) | 0.79 | 0.0299 |
| Delma tincta group | *Delma pax* | 53 | 0.90 | (0.82-0.99) | 0.90 | 0.0245 |
| Lialis group | *Lialis burtonis* (wet biome) | 136 | 0.86 | (0.81-0.91) | 0.91 | 0.0000 |
| Lialis group | *Lialis burtonis* (dry biome) | 142 | 0.78 | (0.73-0.83) | 0.88 | 0.0000 |
| Lialis group | *Lialis jicari* | 96 | 0.79 | (0.72-0.85) | 0.84 | 0.0000 |
| Pygopus lepidopodus group | *Pygopus lepidopodus* (wet biome) | 20 | 0.84 | (0.74-0.93) | 0.95 | 0.0018 |
| Pygopus lepidopodus group | *Pygopus lepidopodus* (dry biome) | 131 | 0.80 | (0.76-0.85) | 0.92 | 0.0000 |
| Pygopus nigriceps group | *Pygopus nigriceps* (wet biome) | 15 | 0.84 | (0.70-0.98) | 0.93 | 0.0256 |
| Pygopus nigriceps group | *Pygopus nigriceps* (dry biome) | 69 | 0.81 | (0.73-0.90) | 0.85 | 0.0000 |
| Pygopus nigriceps group | *Pygopus schraderi* (wet biome) | 35 | 0.90 | (0.80-1.00) | 0.91 | 0.0404 |
| Pygopus nigriceps group | *Pygopus schraderi* (dry biome) | 51 | 0.75 | (0.69-0.82) | 0.92 | 0.0000 |
| Pygopus nigriceps group | *Pygopus steelescotti* | 39 | 0.79 | (0.70-0.87) | 0.91 | 0.0000 |

Table S4. Scaling patterns for pairs of ontogenetic trajectories in the *Aprasia repens* group. All scaling patterns were obtained from Supplementary Figure S1 by using maximum $SVL$ as the measure of body size. GS = geometric similarity; EA = evolutionary allometry; X = one or both OTUs had insufficient data (i.e., missing or regenerated tails); * indicates that the left OTU has a larger maximum $\overline{BW}$ than the right OTU but both OTUs have the same maximum $SVL$; ** indicates that the right OTU has a larger maximum $\overline{BW}$ than the left OTU but the left OTU has a longer maximum $SVL$ than the right OTU; *** indicates that the left OTU has a longer maximum $SVL$ than the right OTU but both OTUs have the same maximum $\overline{BW}$. Scaling relationships for *A. repens* vs. *A. rostrata* pairs could not be determined because both OTUs had the same maximum $SVL$. See Materials and Methods for more details and Supplementary Data for maximum $SVL$ and $\overline{BW}$.

| Larger Max SVL & BW | Smaller Max SVL & BW | lnSVL vs. lnTail | lnBW vs. lnTail | lnSVL vs. lnBW |
|---|---|---|---|---|
| *Aprasia repens* | *Aprasia litorea* | GS (all ages) | GS (all ages) | GS (all ages) |
| *Aprasia rostrata* | *Aprasia litorea* | EA overlap (slopes equal) | EA overlap (slopes equal) | GS (all ages) |
| *Aprasia repens** | *Aprasia rostrata* | ambiguous | ambiguous | ambiguous |
| *Aprasia repens* | *Aprasia clairae* | X | X | GS (adults) |
| *Aprasia repens* | *Aprasia haroldi* | X | X | GS (adults) |
| *Aprasia rostrata* | *Aprasia clairae* | X | X | EA overlap (slopes equal) |
| *Aprasia rostrata* | *Aprasia haroldi* | X | X | EA overlap (slopes equal) |
| *Aprasia smithi* | *Aprasia clairae* | X | X | GS (all ages) |
| *Aprasia smithi* | *Aprasia litorea* | X | X | GS (all ages) |
| *Aprasia smithi* | *Aprasia haroldi* | X | X | GS (all ages) |
| *Aprasia smithi* | *Aprasia rostrata* | X | X | GS (all ages) |
| *Aprasia clairae* | *Aprasia haroldi*** | X | X | EA overlap (slopes equal) |
| *Aprasia litorea* | *Aprasia haroldi*** | X | X | EA transposition (all ages) |
| *Aprasia smithi* | *Aprasia repens*** | X | X | EA overlap (slopes equal) |
| *Aprasia litorea**** | *Aprasia clairae* | X | X | EA transposition (all ages) |

Table S5. Scaling patterns for pairs of ontogenetic trajectories in the southeastern Australia *Aprasia* group. All scaling patterns were obtained from Supplementary Figure S2 by using maximum $SVL$ as the measure of body size. GS = geometric similarity; EA = evolutionary allometry; ** indicates that the right OTU has a larger maximum $\overline{BW}$ than the left OTU but the left OTU has a longer maximum $SVL$ than the right OTU. See Materials and Methods for more details and Supplementary Data for maximum $SVL$ and $\overline{BW}$.

| **Larger Max SVL & BW** | **Smaller Max SVL & BW** | **lnSVL vs. lnTail** | **lnBW vs. lnTail** | **lnSVL vs. lnBW** |
|---|---|---|---|---|
| *Aprasia parapulchella* | *Aprasia inaurita* | EA transposition (all ages) | EA overlap (slopes not equal) | GS (all ages) |
| *Aprasia pseudopulchella* | *Aprasia inaurita* | EA overlap (slopes equal) | EA transposition (juveniles) | EA transposition (all ages) |
| *Aprasia pseudopulchella* | *Aprasia parapulchella*** | GS (all ages) | EA overlap (slopes equal) | EA transposition (all ages) |
| *Aprasia parapulchella* | *Aprasia striolata*** | EA transposition (all ages) | EA transposition (all ages) | EA transposition (all ages) |
| *Aprasia pseudopulchella* | *Aprasia striolata*** | EA transposition (all ages) | EA transposition (all ages) | EA transposition (all ages) |
| *Aprasia inaurita* | *Aprasia striolata*** | EA transposition (all ages) | EA transposition (adults) | EA transposition (adults) |

Table S6. Scaling patterns for pairs of ontogenetic trajectories in the *Delma australis*, *D. fraseri*, and *D. impar* groups. All scaling patterns were obtained from Supplementary Figure S3 by using maximum $SVL$ as the measure of body size. GS = geometric similarity; EA = evolutionary allometry; * indicates that the left OTU has a larger maximum $\overline{BW}$ than the right OTU but both OTUs have the same maximum $SVL$; ** indicates that the right OTU has a larger maximum $\overline{BW}$ than the left OTU but the left OTU has a longer maximum $SVL$ than the right OTU; *** indicates that the left OTU has a longer maximum $SVL$ than the right OTU but both OTUs have the same maximum $\overline{BW}$. Scaling relationships for *D. petersoni* vs. *D. fraseri* pairs could not be determined because both OTUs had the same maximum $SVL$. See Materials and Methods for more details and Supplementary Data for maximum $SVL$ and $\overline{BW}$.

| **Larger Max SVL & BW** | **Smaller Max SVL & BW** | **lnSVL vs. lnTail** | **lnBW vs. lnTail** | **lnSVL vs. lnBW** |
|---|---|---|---|---|
| *Delma australis**** | *Delma hebesa* | EA overlap (slopes equal) | EA overlap (slopes not equal) | EA overlap (slopes not equal) |
| *Delma petersoni** | *Delma fraseri* | ambiguous | ambiguous | ambiguous |
| *Delma molleri* | *Delma impar*** | EA transposition (all ages) | EA transposition (all ages) | EA overlap (slopes equal) |
| *Delma plebeia* | *Delma impar*** | EA transposition (all ages) | EA transposition (adults) | EA overlap (slopes not equal) |
| *Delma plebeia* | *Delma molleri*** | EA overlap (slopes equal) | EA overlap (slopes equal) | EA overlap (slopes not equal) |

Table S7. Scaling patterns for pairs of ontogenetic trajectories in the *Delma nasuta* group. All scaling patterns were obtained from Supplementary Figure S4 by using maximum $SVL$ as the measure of body size. GS = geometric similarity; EA = evolutionary allometry; ** indicates that the right OTU has a larger maximum $\overline{BW}$ than the left OTU but the left OTU has a longer maximum $SV$L than the right OTU. See Materials and Methods for more details and Supplementary Data for maximum $SVL$ and $\overline{BW}$.

| Larger Max SVL & BW | Smaller Max SVL & BW | lnSVL vs. lnTail | lnBW vs. lnTail | lnSVL vs. lnBW |
|---|---|---|---|---|
| *Delma butleri* | *Delma haroldi* | GS (all ages) | GS (all ages) | GS (all ages) |
| *Delma grayii* | *Delma butleri* | EA transposition (adults) | EA transposition (all ages) | EA transposition (all ages) |
| *Delma grayii* | *Delma haroldi* | EA overlap (slopes equal) | EA overlap (slopes equal) | EA overlap (slopes equal) |
| *Delma inornata* | *Delma butleri* | GS (juveniles) | EA overlap (slopes equal) | EA transposition (all ages) |
| *Delma inornata* | *Delma grayii* | GS (all ages) | GS (all ages) | EA transposition (all ages) |
| *Delma inornata* | *Delma haroldi* | GS (all ages) | GS (all ages) | EA transposition (all ages) |
| *Delma inornata* | *Delma nasuta* | GS (all ages) | GS (all ages) | EA transposition (all ages) |
| *Delma nasuta* | *Delma butleri* | EA transposition (adults) | EA transposition (all ages) | EA overlap (slopes equal) |
| *Delma nasuta* | *Delma haroldi* | GS (all ages) | GS (all ages) | EA overlap (slopes equal) |
| *Delma grayii* | *Delma nasuta*** | EA transposition (all ages) | EA transposition (all ages) | EA overlap (slopes equal) |

Table S8. Scaling patterns for pairs of ontogenetic trajectories in the *Delma tincta* group. All scaling patterns were obtained from Supplementary Figure S5 by using maximum $SVL$ as the measure of body size. GS = geometric similarity; EA = evolutionary allometry; * indicates that the left OTU has a larger maximum $\overline{BW}$ than the right OTU but both OTUs have the same maximum $SVL$. Scaling relationships for *D. borea* vs. *D. pax* pairs could not be determined because both OTUs had the same maximum $SVL$. See Materials and Methods for more details and Supplementary Data for maximum $SVL$ and $\overline{BW}$.

| Larger Max SVL & BW | Smaller Max SVL & BW | lnSVL vs. lnTail | lnBW vs. lnTail | lnSVL vs. lnBW |
|---|---|---|---|---|
| *Delma borea* | *Delma desmosa* | GS (all ages) | EA overlap (slopes not equal) | EA overlap (slopes equal) |
| *Delma borea** | *Delma pax* | ambiguous | ambiguous | ambiguous |
| *Delma pax* | *Delma desmosa* | GS (all ages) | GS (all ages) | GS (all ages) |

Table S9. Scaling patterns for pairs of ontogenetic trajectories in the *Lialis* and *Pygopus lepidopodus* groups. All scaling patterns were obtained from Supplementary Figure S6 by using maximum $SVL$ as the measure of body size. GS = geometric similarity; EA = evolutionary allometry; ** indicates that the right OTU has a larger maximum $\overline{BW}$ than the left OTU but the left OTU has a longer maximum $SVL$ than the right OTU. See Materials and Methods for more details and Supplementary Data for maximum $SVL$ and $\overline{BW}$.

| **Larger Max SVL & BW** | **Smaller Max SVL & BW** | **lnSVL vs. lnTail** | **lnBW vs. lnTail** | **lnSVL vs. lnBW** |
|---|---|---|---|---|
| *Lialis burtonis* (wet biome) | *Lialis burtonis* (dry biome) | GS (all ages) | GS (all ages) | GS (adults) |
| *Lialis burtonis* (wet biome) | *Lialis jicari* | GS (adults) | GS (adults) | GS (all ages) |
| *Lialis jicari* | *Lialis burtonis*** (dry biome) | EA transposition (adults) | EA transposition (all ages) | EA transposition (all ages) |
| *Pygopus lepidopodus* (wet biome) | *Pygopus lepidopodus* (dry biome) | GS (all ages) | GS (all ages) | GS (all ages) |

Table S10. Scaling patterns for pairs of ontogenetic trajectories in the *Pygopus nigriceps* group. All scaling patterns were obtained from Supplementary Figure S7 by using maximum $SVL$ as the measure of body size. GS = geometric similarity; EA = evolutionary allometry; ** indicates that the right OTU has a larger maximum $\overline{BW}$ than the left OTU but the left OTU has a longer maximum $SVL$ than the right OTU. See Materials and Methods for more details and Supplementary Data for maximum $SVL$ and $\overline{BW}$.

| **Larger Max SVL & BW** | **Smaller Max SVL & BW** | **lnSVL vs. lnTail** | **lnBW vs. lnTail** | **lnSVL vs. lnBW** |
|---|---|---|---|---|
| *Pygopus nigriceps* (wet biome) | *Pygopus nigriceps* (dry biome) | GS (adults) | GS (all ages) | GS (all ages) |
| *Pygopus nigriceps* (wet biome) | *Pygopus schraderi* (dry biome) | GS (all ages) | EA transposition (all ages) | EA transposition (all ages) |
| *Pygopus nigriceps* (wet biome) | *Pygopus steelescotti* | EA overlap (slopes not equal) | EA transposition (juveniles) | EA transposition (all ages) |
| *Pygopus schraderi* (wet biome) | *Pygopus schraderi* (dry biome) | GS (all ages) | GS (all ages) | GS (adults) |
| *Pygopus schraderi* (wet biome) | *Pygopus steelescotti* | EA overlap (slopes equal) | GS (adults) | GS (all ages) |
| *Pygopus nigriceps* (wet biome) | *Pygopus schraderi*** (wet biome) | EA overlap (slopes not equal) | EA transposition (all ages) | EA transposition (all ages) |
| *Pygopus nigriceps* (dry biome) | *Pygopus schraderi*** (wet biome) | EA overlap (slopes equal) | EA transposition (all ages) | EA transposition (all ages) |
| *Pygopus nigriceps* (dry biome) | *Pygopus schraderi*** (dry biome) | GS (all ages) | EA transposition (all ages) | EA transposition (all ages) |
| *Pygopus nigriceps* (dry biome) | *Pygopus steelescotti*** | EA overlap (slopes equal) | EA transposition (juveniles) | EA transposition (all ages) |
| *Pygopus steelescotti* | *Pygopus schraderi*** (dry biome) | GS (all ages) | EA overlap (slopes equal) | EA transposition (all ages) |

Table S11. Scaling patterns for pairs of static adult growth trajectories in four species of Darwin's finches belonging to the genus *Geospiza*. Scaling patterns were obtained from log-log plots of geometric mean regressions for logBLG on logWGT, logBDT on logWGT, and logBWD on logWGT relationships shown in Figure 3 of Boag (1984). Each scaling pattern was classified as reflecting either geometric similarity or evolutionary allometry based on the spatial arrangement of the trajectories. Note that nine of the comparisons involved pairs of species while one comparison involved two allopatric populations of *G. fortis*. OTU = operational taxonomic unit; BLG = bill length; BDT = bill depth; BWD = bill width; and WGT = body mass (see Boag 1984 for more details).

| **larger OTU/smaller OTU** | **BLG on WGT** | **BDT on WGT** | **BWD on WGT** |
|---|---|---|---|
| *G. magnirostris* /*G. fortis* (Daphne Major Island) | geometric similarity | evolutionary allometry | evolutionary allometry |
| *G. magnirostris* /*G. fortis* (Santa Cruz Island) | evolutionary allometry | evolutionary allometry | evolutionary allometry |
| *G. magnirostris* /*G. scandens* | geometric similarity | evolutionary allometry | evolutionary allometry |
| *G. magnirostris* /*G. fuliginosa* | evolutionary allometry | evolutionary allometry | evolutionary allometry |
| *G. fortis* (Santa Cruz Island)/*G. fortis* (Daphne Major Island) | geometric similarity | geometric similarity | geometric similarity |
| *G. fortis* (Daphne Major Island)/*G. fuliginosa* | evolutionary allometry | evolutionary allometry | evolutionary allometry |
| *G. scandens* /*G. fortis* (Daphne Major Island) | evolutionary allometry | geometric similarity | geometric similarity |
| *G. scandens* /*G. fuliginosa* | evolutionary allometry | evolutionary allometry | evolutionary allometry |
| *G. fortis* (Santa Cruz Island)/*G. fuliginosa* | evolutionary allometry | evolutionary allometry | evolutionary allometry |
| *G. fortis* (Santa Cruz Island)/*G. scandens* | geometric similarity | evolutionary allometry | evolutionary allometry |
